# DiMeLo-seq: a long-read, single-molecule method for mapping protein-DNA interactions genome-wide

**DOI:** 10.1101/2021.07.06.451383

**Authors:** Nicolas Altemose, Annie Maslan, Owen K. Smith, Kousik Sundararajan, Rachel R. Brown, Angela M. Detweiler, Norma Neff, Karen H. Miga, Aaron F. Straight, Aaron Streets

**Affiliations:** Department of Bioengineering, University of California, Berkeley, CA 94720; UC Berkeley-UCSF Graduate Program in Bioengineering, University of California, Berkeley, Berkeley, CA 94720; Center for Computational Biology, University of California, Berkeley, CA 94720; Department of Biochemistry, Stanford University, Stanford, CA 94305; Department of Chemical and Systems Biology, Stanford University, Stanford, CA 94305; Chan Zuckerberg Biohub, San Francisco, CA 94158; Department of Molecular & Cell Biology, University of California, Santa Cruz, CA 95064; UC Santa Cruz Genomics Institute, University of California, Santa Cruz, CA 95064; Department of Molecular & Cell Biology, University of California, Berkeley, CA 94720

**Author notes:** These authors co-supervised the study; to whom correspondence should be addressed. These authors contributed equally (Nicolas Altemose, Annie Maslan, Owen K. Smith, Kousik Sundararajan).

## Abstract

Molecular studies of genome regulation often rely on the ability to map where specific proteins interact with genomic DNA. Existing techniques for mapping protein-DNA interactions genome-wide rely on DNA amplification methods followed by sequencing with short reads, which dissociates joint binding information at neighboring sites, removes endogenous DNA methylation information, and precludes the ability to reliably map interactions in repetitive regions of the genome. To address these limitations, we created a new protein-DNA mapping method, called **D**irected **M**ethylation with **L**ong-read **seq**uencing (DiMeLo-seq), which methylates DNA near each target protein’s DNA binding site *in situ*, then leverages the ability to distinguish methylated and unmethylated bases on long, native DNA molecules using long-read, single-molecule sequencing technologies. We demonstrate the optimization and utility of this method by mapping the interaction sites of a variety of different proteins and histone modifications across the human genome, achieving a single-molecule binding site resolution of less than 200 bp. Furthermore, we mapped the positions of the centromeric histone H3 variant CENP-A in repetitive regions that are unmappable with short reads, while simultaneously analyzing endogenous CpG methylation and joint binding events on single molecules. DiMeLo-seq is a versatile method that can provide multimodal and truly genome-wide information for investigating protein-DNA interactions.

## Introduction

Genomic DNA must be decoded and maintained by proteins that read, regulate, replicate, recombine, and repair it. Mapping where and how these proteins interact with DNA can provide key insights into how they function or malfunction in healthy and diseased cells. Several powerful approaches have been developed to map where a specific protein interacts with DNA genome-wide, including DamID, ChIP-seq, CUT&RUN, and related methods (van Steensel and Henikoff 2000; Mikkelsen et al. 2007; Robertson et al. 2007; Johnson et al. 2007; Barski et al. 2007; Skene and Henikoff 2017). These approaches involve selectively amplifying short DNA fragments from regions bound by a particular protein of interest, determining the sequence of those DNA molecules using high-throughput DNA sequencing, and then mapping those sequences back to a reference genome, using sequencing coverage as a measure of protein-DNA interaction frequency. While these methods have proven to be extremely useful for studying DNA-binding proteins and chromatin modifications (Rivera and Ren 2013), they rely on strategies for enrichment and detection that present important limitations.

Firstly, the process of DNA amplification fails to copy DNA modification information, like endogenous CpG methylation, from the native DNA molecules to the amplified and sequenced library DNA. This prevents simultaneous measurement of protein-DNA interactions and DNA methylation and limits the amount of information that can be gleaned about the relationship between these regulatory elements. Secondly, amplification-based enrichment typically relies on nonlinear methods like PCR to yield enough amplified product for sequencing. Due to amplification bias intrinsic to PCR, sequencing coverage is only a semi-quantitative readout of protein-DNA interaction frequencies.

Furthermore, these approaches rely on digesting or shearing DNA into short fragments for enrichment and standard high-throughput DNA sequencing for detection, which produces short sequencing reads typically under 250 bp in length. While fragment length defines binding site resolution in these techniques, this dependence on short fragments destroys joint long-range binding information from the same chromatin fibers. Additionally, short sequencing reads cannot be reliably mapped throughout highly repetitive regions of the genome. Repetitive regions of the human genome have provided a major challenge for genome assembly and mapping methods due to the difficulty of unambiguously assigning short DNA sequencing reads to their unique positions in the genome. Recent efforts using long-read sequencing technologies have, for the first time, provided a complete assembly of repetitive regions across the human genome (Nurk et al. 2021). However, unambiguously mapping short-read sequencing data remains impossible in many repetitive genomic regions, limiting our ability to address lingering biological questions about the roles of repeat sequences in cell division, protein synthesis, aging, and genome regulation.

These limitations underline the need for new protein-DNA interaction mapping methods that fully leverage the power of long-read, single-molecule sequencing technologies, including their ability to read out DNA modifications directly. To address this need, we developed Directed Methylation with Long-read sequencing (DiMeLo-seq; from *dímelo*, pronounced DEE-meh-lo). DiMeLo-seq provides the ability to map protein-DNA interactions with high resolution on native, long, single, sequenced DNA molecules, while simultaneously measuring endogenous DNA modifications and sequence variation. Each of these features provides an opportunity to study genome regulation in unprecedented ways. Recent technologies have begun to take advantage of long-read sequencing to identify accessible regions and CpG methylation on native single molecules, but they cannot target specific protein-DNA interactions (Abdulhay et al. 2020; Stergachis et al. 2020; Lee et al. 2020; Shipony et al. 2020; Wang et al. 2019). Here we extend these capabilities to map specific regulatory elements and demonstrate the advantages of DiMeLo-Seq by mapping lamina associated domains, CTCF binding sites, histone modifications/variants, and CpG methylation across the genome and through complex repetitive domains.

## Results

### 1. *Antibody-directed DNA adenine methylation enables histone-specific DNA methylation of chromatin* in vitro

DiMeLo-seq combines elements of antibody-directed protein-DNA mapping approaches (Skene and Henikoff 2017; Schmid, Durussel, and Laemmli 2004; van Schaik et al. 2020) to deposit methylation marks near a specific target protein, then uses long-read sequencing to read out these exogenous methylation marks directly (Abdulhay et al. 2020; Stergachis et al. 2020; Lee et al. 2020; Shipony et al. 2020; Wang et al. 2019). Taking advantage of the lack of N^6^-methyl-deoxyadenosine in human DNA (O’Brown et al. 2019), we fused the antibody-binding Protein A to the nonspecific deoxyadenosine methyltransferase Hia5 (Stergachis et al. 2020; Drozdz et al. 2012) (pA-Hia5) to catalyze the formation of N^6^-methyl-deoxyadenosine (hereafter mA) in the DNA proximal to targeted chromatin-associated proteins (Fig. 1a). First, nuclei are permeabilized, primary antibodies are bound to the protein of interest, and any unbound antibody is washed away. Then, pA-Hia5 is bound to the antibody, and any unbound pA-Hia5 is washed away. The nuclei are then incubated in a buffer containing the methyl donor S-adenosyl methionine (SAM) to activate adenine methylation in the vicinity of the protein of interest (van Schaik et al. 2020). Finally, genomic DNA is isolated and sequenced using modification-sensitive, long-read sequencing (Fig. 1a) with mA basecalls providing a readout of the sites of DNA-protein interactions. This approach provides a distinct advantage in the ability to detect multiple protein binding events on a long single DNA molecule, which would not be possible with short-read sequencing (Fig. 1b). This protocol avoids amplification biases, allowing for identification of modifications that are proportional to the frequency of a given molecular interaction in cells (Fig. 1c). Modification-sensitive readout allows for the simultaneous detection of both exogenous antibody-directed adenine methylation and endogenous CpG methylation on single molecules (Fig. 1d). Finally, using long-read sequencing to identify specific protein-DNA interactions allows for improvements in mappability within highly repetitive regions of the genome (Fig. 1e; full workflow in Supplementary Fig. 1). Overall, these improvements allow investigation of protein-DNA interactions on the single-molecule level, including in challenging genomic regions, with resolution and specificity that was not previously possible.

**Figure 1.**
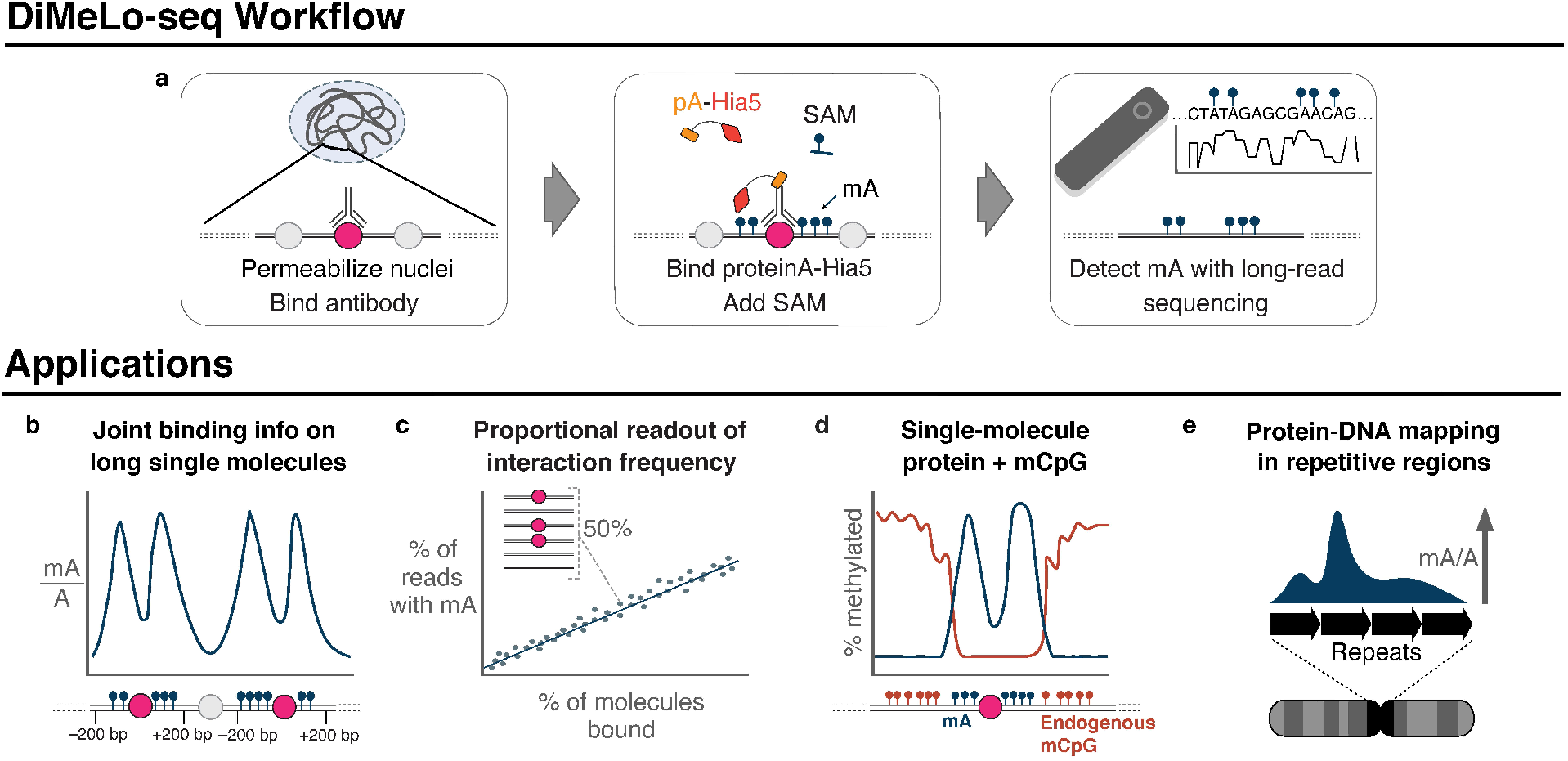
High-resolution, genome-wide mapping of DNA-protein interactions with DiMeLo-seq. **a**, Schematic of the DiMeLo-seq workflow for the mapping of DNA-protein interactions. **b**, DiMeLo-seq can be used to map joint binding info of proteins on long, single molecules. **c**, As an amplification-free method, the readout of DiMeLo-seq is proportional to the DNA-protein interaction frequency in cells. **d**, This method can be multiplexed to map both single-molecule DNA-protein interactions and endogenous CpG methylation. **e**, DNA-protein interactions can be mapped across the entire genome, including to repetitive regions that were previously challenging to map.

We expressed and purified recombinant pA-Hia5 and tested its methylation activity on purified DNA using the methylation-sensitive restriction enzyme DpnI, which only cuts GATC sites when adenine is methylated. Increasing concentrations of Hia5, pA-Hia5, or Protein A/G-Hia5 (pAG-Hia5) in the presence of SAM caused the DNA to become sensitive to DpnI digestion, confirming the methyltransferase activity of the purified fusion proteins (Supplementary Fig. 2a,b). We tested the ability of pA-Hia5 to methylate accessible regions of DNA on *in vitro* reconstituted chromatin assembled on an 18x array of the 601 nucleosome positioning sequence (Lowary and Widom 1998). Co-incubation of chromatin together with free pA-Hia5 and SAM followed by long-read sequencing and methylation-sensitive basecalling resulted in structured patterns of oligonucleosome footprinting (Supplementary Fig. 2c-e), as reported previously for reconstituted chromatin incubated with another exogenous methyltransferase, EcoGII (Abdulhay et al. 2020).

To test the ability of antibodies to direct the methyltransferase activity of pA-Hia5, we incubated an antibody targeting the histone H3 variant CENP-A with chromatin that was reconstituted using either CENP-A or canonical histone H3 containing nucleosomes. We conjugated the chromatin arrays to streptavidin-coated magnetic beads by biotinylating the DNA ends, thereby allowing us to wash away unbound antibody and pA-Hia5 prior to activating methylation with SAM (Fig. 2a). Following activation, we immunostained chromatin-conjugated beads with an anti-mA antibody and demonstrated a significant increase in mA immunofluorescence signal only when CENP-A chromatin, but not H3 chromatin, was incubated with pA-Hia5 following CENP-A antibody binding (Fig. 2b-c, Supplementary Fig. 2f), suggesting specific methylation by antibody-directed pA-Hia5. Long-read sequencing detected mA on DNA after directed methylation, albeit to a lower extent when compared to incubation with free pA-Hia5 (Fig. 2c). On average, CENP-A-directed methylation of CENP-A chromatin appeared depleted at the central axis of the nucleosome where the 601 sequence positions the nucleosome dyad (Fig. 2d). On individual reads, we observed protection patterns that span between 100 - 170 bp, consistent with nucleosome occupancy protecting the DNA from antibody-directed methylation (Fig. 2d,e), similar to the free pA-Hia5 condition (Supplementary Fig. 2d,e). The center of these footprints align within ~ 30 bp of the 601 dyad position (Fig. 2f,g). We determined the extent of methylation in the vicinity of nucleosome-sized footprints on sparsely occupied chromatin after computationally filtering for nucleosomes spaced apart by at least 1 601 position. In contrast to the free pA-Hia5 condition, for which we observed a high prevalence of methylation on any region not protected by nucleosomes, in the antibody-directed pA-Hia5 condition, the average probability of methylation rapidly decreased in the vicinity of the nucleosome footprints (within ~75 bp, Fig. 2h), suggesting preferential methylation of deoxyadenosine closest to the antibody-bound nucleosome. We observed similar results for H3-antibody-directed methylation using pAG-Hia5 (Supplementary Fig. 2h-j). We conclude that directing pA-Hia5 activity using a histone-specific antibody targets specific methylation in proximity to the nucleosome of interest *in vitro*.

**Figure 2.**
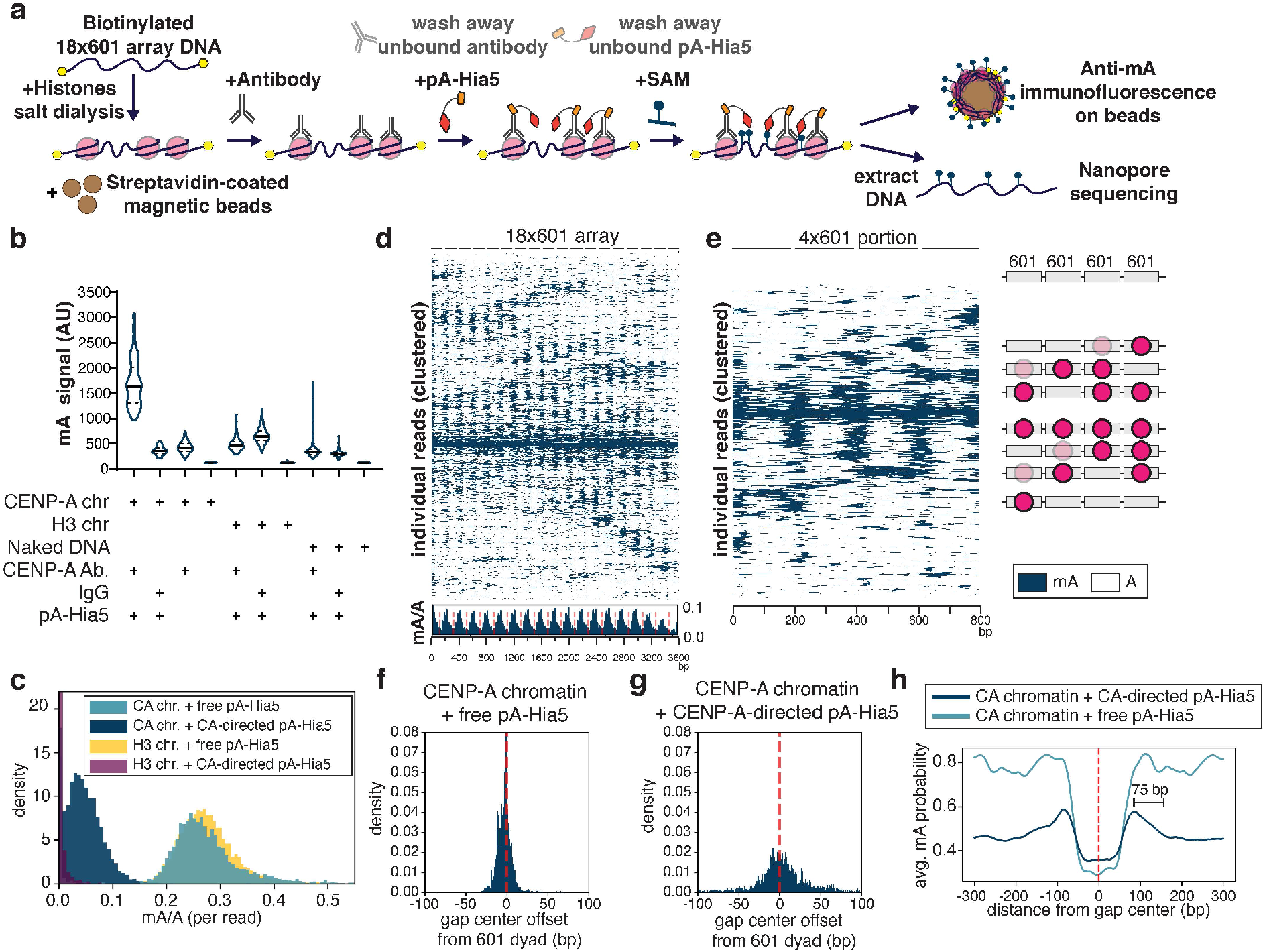
Antibody directed methylation of artificial chromatin and long-read sequencing. **a**, Schematic of directed methylation of artificial chromatin depicting biotinylated chromatin reconstitution, specific antibody binding, pA-Hia5 targeting, SAM addition and activation, followed by long-read sequencing of methylated DNA extracted from chromatin. **b**, Immunofluorescence signal on chromatin-coated beads following antibody-directed methylation. Solid line - median, dashed line - quartiles. n > 90 beads/condition. **c**, Histogram of fraction of methylation (mA/A) on reads from CENP-A or H3 chromatin methylated with free pA-Hia5 or CENP-A-directed pA-Hia5. **d,e**, Heatmap showing methylation on 2000 individual reads from CENP-A chromatin methylation with CENP-A-directed pA-Hia5, hierarchically clustered by jaccard distance over the entire 18×601 array (d) or a subset 4×601 region along with cartoons depicting predicted nucleosome positions (e). Inset in d. shows average mA/A on every base position of 18×601 array. (red dashed line indicates 601 dyad position). **f,g**, Position of centers of 100-180 bp gaps in mA signal relative to the closest theoretical 601 dyad position (red dashed line) on reads from free pA-Hia5 condition (f) or CENP-A-directed pA-Hia5 condition (g). **h**, Average methylation probability score (from Megalodon base-calling) near the center of 100-180 bp gaps (red dashed line) in mA signal on reads from free pA-Hia5 condition or CENP-A directed pA-Hia5 condition.

### 2. *Optimization of LMNB1 mapping* in situ

We next optimized DiMeLo-seq for mapping protein-DNA interactions *in situ* in permeabilized nuclei from a human cell line (HEK293T). To do this, we mapped the interaction sites of Lamin B1 (LMNB1), which is a long, filamentous protein that forms part of the nuclear lamina on the inside of the nuclear envelope, and which is often targeted in DamID studies to profile lamina associated domains (LADs) (Guelen et al. 2008). Large regions of the genome that are almost always in contact with the nuclear lamina across cell types are called constitutive lamina associated domains (cLADs), and these tend to be heterochromatic, gene-poor, and transcriptionally quiet. Regions that are almost never in contact with the nuclear lamina across cell types and instead reside in the nuclear interior are called constitutive inter-LADs (ciLADs), and they tend to be euchromatic, gene-rich, and transcriptionally active (Fig. 3a). Other regions can vary in their lamina contact frequency between cell types and/or between cells of the same type. We chose LMNB1 as an initial target because (i) cLADs and ciLADs provide well-characterized on-target and off-target control regions, respectively; (ii) LMNB1 has a very large binding footprint (median LAD size is 500 kb), meaning we can detect its interactions even with very low sequencing coverage; (iii) LMNB1 localization at the nuclear lamina can be easily visualized by immunofluorescence, allowing us to use microscopy for intermediate quality control during each step of the protocol (Supplementary Fig. 3); and (iv) we have previously generated LMNB1 DamID data from HEK293T cells using bulk and single-cell protocols, providing ample reference materials (Altemose et al. 2020).

**Figure 3.**
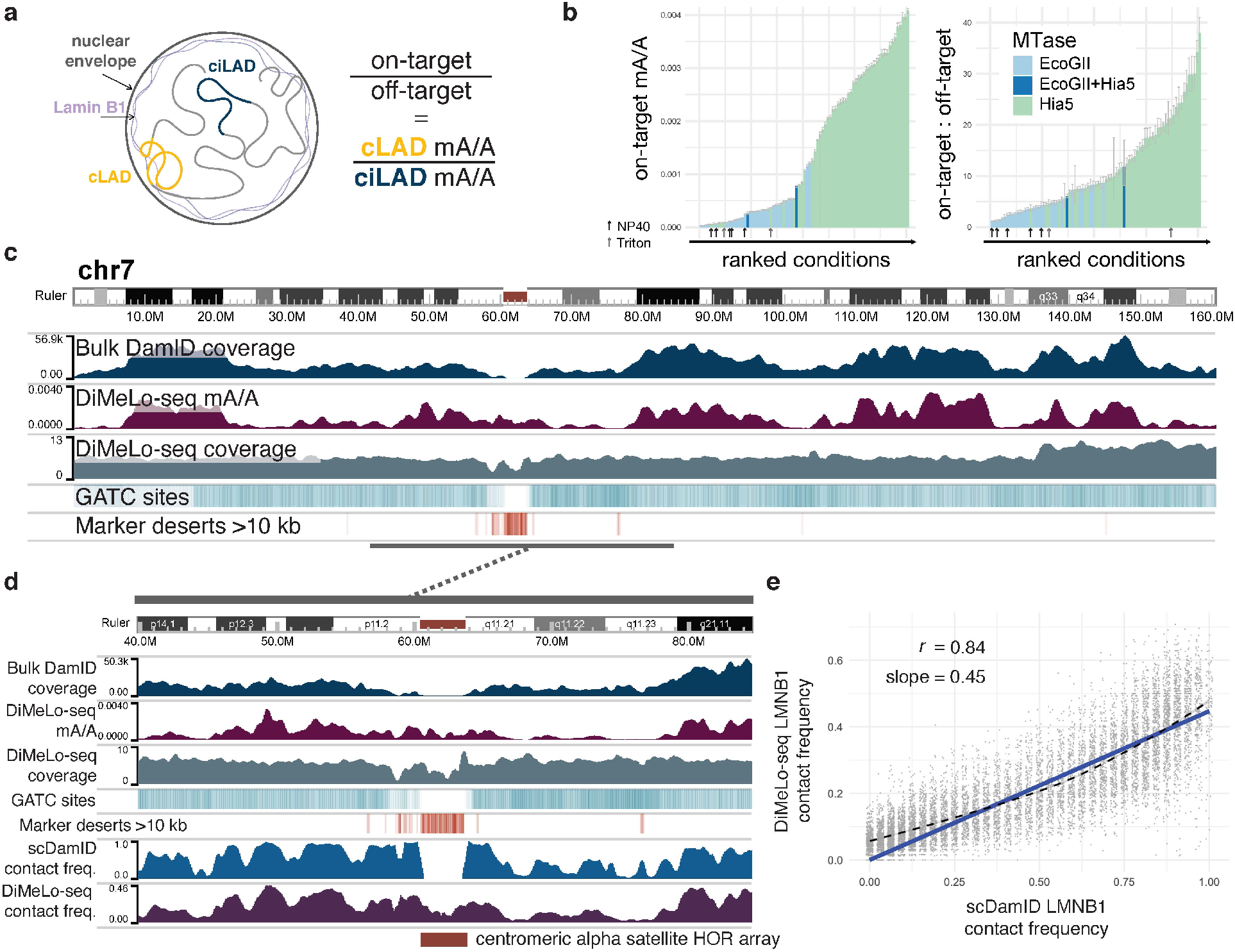
Optimization of DiMeLo-seq targeting Lamin B1 *in situ*. **a**, Schematic of interactions between LMNB1 and lamina-associated domains, and the use of mA levels in cLADs and ciLADs to estimate on-target and off-target mA. **b**, Barplots showing for all protocol conditions tested the proportion of all A bases (q>10) on all reads called as methylated (p>0.9; abbreviated mA/A) in on-target (cLAD) regions (left plot) and the ratio of these mA levels compared to off-target (ciLAD) regions (right plot). Bars are ordered by increasing height and colored by the methyltransferase used, showing improved performance with Hia5. Samples for which NP40 or Triton X-100 were used for permeabilization instead of digitonin are indicated with black and gray arrows, respectively. Complete data are in Supplementary Table 1. **c**, A browser image across all of chromosome 7 comparing *in situ* LMNB1-targeted DiMeLo-seq to *in vivo* LMNB1-tethered DamID data (blue) (Altemose et al. 2020). The coverage of each region by simulated DpnI digestion fragments (splitting reference at GATC sites) between 150 and 750 bp (sequenceable range) is indicated by a teal heatmap track (range 0 to 0.7). The presence of intervals longer than 10 kb between unique 51-mers in the reference, a measure of mappability, is indicated with an orange heatmap track. **d**, A closer view of the centromere on chr7, with added tracks at the bottom illustrating LMNB1 contact frequencies from single-cell DamID data (Altemose et al. 2020), as well as from DiMeLo-seq data (Methods). **e**, For a quality-filtered set of 100 kb bins genome-wide (Methods), a comparison of scDamID and DiMeLo-seq contact frequency estimates. A zero-intercept linear regression line (blue) and loess line (dashed black) are overlaid.

To assess the performance of the LMNB1-targeted DiMeLo-seq protocol, we quantified the proportion of adenines that were called as methylated across all reads mapping to cLADs (on-target regions), and across all reads mapping to ciLADs (off-target regions). We evaluated the performance of each iteration of the protocol using both the on-target methylation rate (as a proxy for sensitivity) and the on-target:off-target ratio (as a proxy for signal-to-background), aiming to increase both. We found that we could reliably estimate these values using only ~0.2X genome-wide coverage per sample, allowing us to multiplex multiple conditions on the same MinION flow cell and achieve sufficient coverage after only 24 hours of sequencing. We developed a rapid pipeline for testing variations of many components of the protocol, allowing us to go from harvested cells to fully analyzed data in under 60 hours (Methods). With this optimization pipeline, we tested 80 different conditions (Fig. 3b), varying the following: methyltransferase type (Hia5 vs EcoGII), input cell numbers, detergents, primary antibody concentrations, the use of secondary antibodies, enzyme concentrations, incubation temperatures, methylation incubation times, methylation buffers, and methyl donor concentrations (Methods). Our final optimized protocol is available at protocols.io (link in Methods). Using the finalized protocol and applying a stringent methylation probability threshold of 0.9 (Supplementary Fig. 3, Methods), we regularly achieve on-target methylation of 0.2-0.4% of adenines in cLADs, with an on-target:off-target ratio in the range of 15-30 (Supplementary Table 1).

We also verified that there is very little loss of performance when using cryopreserved or lightly fixed cells, when using between 1-5 million cells per replicate, or when using concanavalin-A coated magnetic beads to carry out cell washing steps by magnetic separation instead of centrifugation (Methods, Supplementary Table 1). Surprisingly, we found no improvement in on-target methylation when using a secondary antibody to recruit more methyltransferase molecules to each site, perhaps due to steric effects, and we saw no improvement when increasing the linker length between pA and Hia5 (Supplementary Fig. 3 and Supplementary Table 1). We saw a slight drop in performance when using pAG-Hia5 compared to pA-Hia5, also potentially due to steric effects. We also found that cell permeabilization with NP40 or TritonX-100 (vs. standard digitonin) actively reduces methylation downstream (Fig. 3b and Supplementary Table 1). While optimization was carried out in HEK293T cells, we also validated that the protocol worked in other human cell lines as well: Hap1, GM12878, and HG002. Finally, to confirm antibody specificity, we performed IgG isotype controls and free-floating Hia5 controls to measure nonspecific methylation and DNA accessibility, respectively (Methods, Supplementary Table 2).

Unlike other protein-DNA mapping methods, which use sequencing coverage as a readout of interaction frequency, DiMeLo-seq sequences the genome uniformly and extracts a separate layer of information (adenine methylation) to measure protein-DNA interaction frequencies across the genome. Thus, we can plot DiMeLo-seq’s coverage and methylation frequency as separate tracks in a browser representing the T2T-CHM13 complete reference sequence, and we can compare these to the results obtained for the same protein target in the same cells by conventional bulk DamID (Fig. 3c). We found that the two methods are highly concordant in the non-repetitive parts of the genome (Spearman correlation = 0.71 in 1 Mb bins), but conventional DamID achieves little-to-no coverage across pericentromeric regions (Fig. 3c). This is due in part to the low availability of unique sequence markers to map short reads to in the pericentromere, but also to the low frequency of GATC (the binding motif for Dam and DpnI in the DamID protocol) within centromeric repeats (Fig. 3c) (Sobecki et al. 2018). DiMeLo-seq, unlike DamID, produces long reads that can be uniquely mapped across the centromeric region of chromosome 7, revealing that this region has an intermediate level of contact with the nuclear lamina (Fig. 3c,d). Obtaining higher coverage with even longer reads in these regions will allow dissection of these differences at much finer resolution.

Because DiMeLo-seq directly probes unamplified genomic DNA, the frequency of single-molecule protein-DNA interactions should be proportional to interaction frequencies in single cells. We leveraged single-cell LMNB1-tethered DamID data from the same cell line (Altemose et al. 2020) to assess the relationship between DiMeLo-seq methylation and known protein-DNA interaction frequencies. In single-cell DamID, each 100 kb bin of the genome is given a binary classification indicating whether it was in contact with the nuclear lamina or not in that particular cell during an ~18 hour incubation period when Dam-LMNB1 is expressed *in vivo* (Kind et al. 2015). Across a sample of 32 single cells, we used these binary classifications to determine a “contact frequency” (CF) for each bin of the genome across the sample of cells (Kind et al. 2015; Altemose et al. 2020). We then performed a similar binary classification of individual DiMeLo-seq reads based on each read’s proportion of methylated adenines. We found that, among reads with at least 500 adenine basecalls (Methods), detection of even a single mA at a 0.9 probability threshold could identify 51% reads from cLADs (51% sensitivity), while only capturing 6% of ciLADs (94% specificity). We then computed a DiMeLo-seq CF in each 100 kb bin, and we compared this to the scDamID CF in the same bin genome-wide. We filtered to bins with high DiMeLo-seq coverage and confident single-cell classifications (see Methods). This revealed a nearly linear relationship between the two contact frequency estimates, with a Pearson correlation of 0.84 (Fig. 3e). The slope of the best-fit zero-intercept line, 0.45, provides an estimate of the overall single-molecule sensitivity of the method. We note that scDamID tends to slightly overestimate intermediate interaction frequencies, attributable to the *in vivo* vs *in situ* nature of the two protocols (van Schaik et al. 2020). This analysis demonstrates the proportionality between single-molecule interaction frequencies measured by DiMeLo-seq and single-cell variability, confirming that DiMeLo-seq is capable of profiling heterogeneity in protein-DNA interactions at the single-cell level.

### 3. Joint analysis of CTCF binding and CpG methylation on single molecules

DiMeLo-seq measures protein-DNA interactions in the context of the local chromatin environment by simultaneously detecting endogenous CpG methylation, nucleosome occupancy, and protein binding. To highlight this feature of DiMeLo-seq, we targeted CTCF, a protein that strongly positions surrounding nucleosomes and whose binding is inhibited by CpG methylation (Bell and Felsenfeld 2000). We first validated that targeted methylation is specific to CTCF in GM12878 cells by calculating the fraction of adenines that are methylated within GM12878 CTCF ChIP-seq peaks relative to the fraction of adenines methylated outside of these peaks. We chose to target CTCF in GM12878 cells because GM12878 is an ENCODE Tier 1 cell line with abundant ChIP-seq reference datasets. We measured a 22-fold increase in targeted methylation over background in our CTCF-targeted sample (Supplementary Fig. 4a). We also measured a 6-fold mA/A enrichment in the free pA-Hia5 control in CTCF ChIP-seq peaks, which reflects the fact that many CTCF binding sites overlap with accessible regions of the genome where pA-Hia5 can methylate more easily (Song et al. 2011). However, both the free pA-Hia5 and the IgG controls produced significantly less targeted methylation than the CTCF-targeted sample (Supplementary Fig. 4a).

As further validation of DiMeLo-seq’s concordance with ChIP-seq data and to visualize protein binding on single molecules, we analyzed mA and mCpG across individual molecules spanning CTCF motifs within ChIP-seq peaks of various strengths (Fig. 4a). DiMeLo-seq signal tracks with ChIP-seq signal strength, with mA density decreasing from the top to bottom quartiles of ChIP-seq peak signal. We observed an increase in local mA surrounding the binding motif, with a periodic decay in methylation from the peak center, indicating methylation of neighboring linker DNA between strongly positioned nucleosomes (Supplementary Fig. 4b). The 76 bp dip at the center of the binding peak reflects CTCF’s binding footprint (Boyle et al. 2011; Klenova et al. 1993; Lobanenkov et al. 1990) and is evident even on single molecules. Interestingly, we observed an asymmetric methylation profile, with stronger methylation 5’ of the CTCF motif. This increased methylation relative to 3’ of the motif extends beyond the central peak to the neighboring linker DNA. This asymmetry could be explained by antibody positioning and the presence of a neighboring protein, such as cohesin, 3’ to the binding motif. The antibody binds the C terminus of CTCF, thereby positioning pA-Hia5 closer to the 5’ end of the binding motif. In addition, the presence of cohesin 3’ to the binding motif may be sterically hindering tethered pA-Hia5’s ability to methylate DNA (Nora et al. 2020). The free pA-Hia5 control profile supports this hypothesis, as there is a less dramatic asymmetry that does not extend to the neighboring linker DNA; free pA-Hia5 may be similarly blocked by cohesin but is able to freely diffuse around cohesin, unlike pA-Hia5 tethered to CTCF (Supplementary Fig. 5).

**Figure 4.**
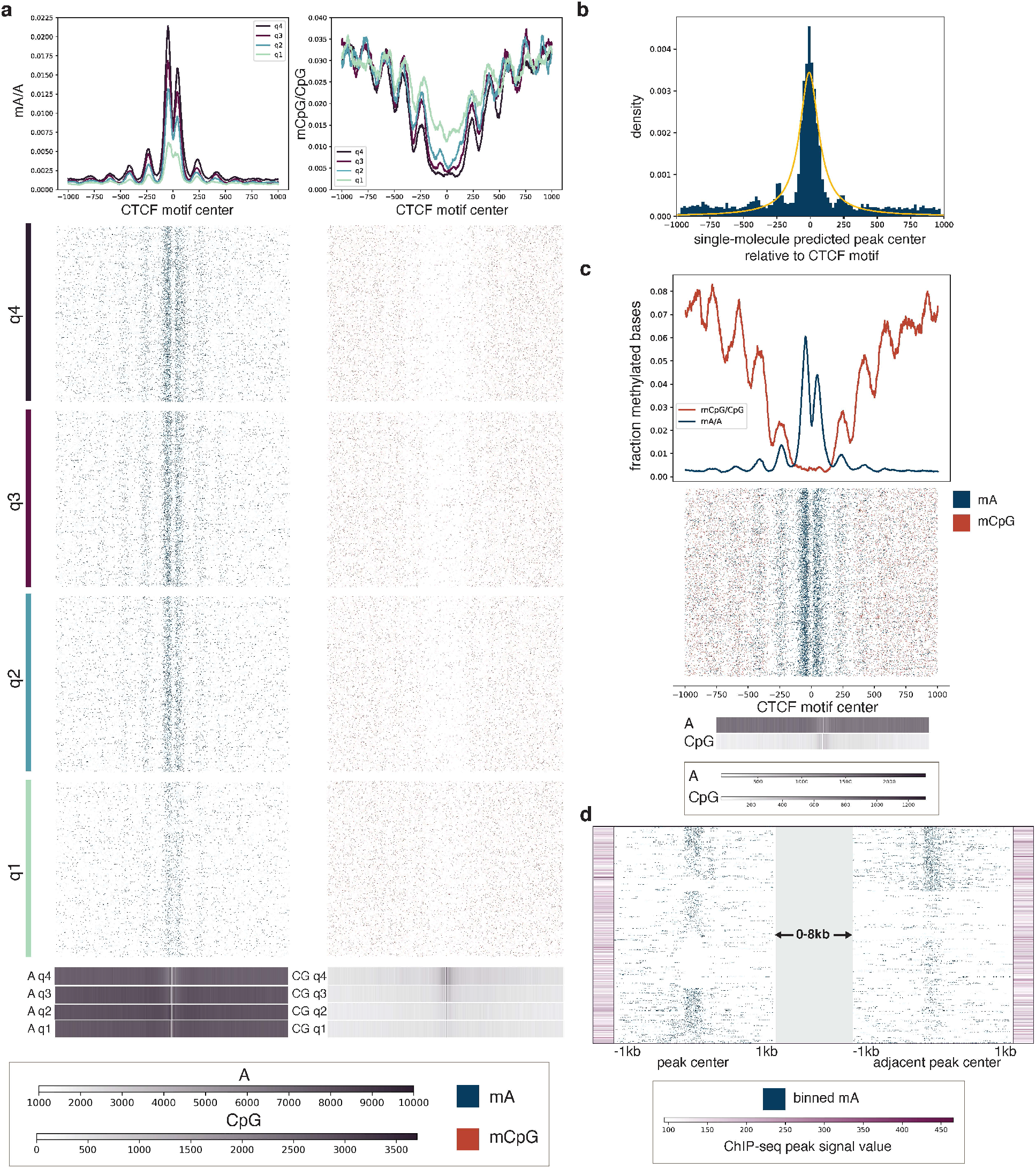
Single-molecule CTCF binding and CpG methylation profiles. **a**, Single molecules spanning CTCF ChIP-seq peaks are shown across quartiles of ChIP-seq peak strength within 1 kb of the peak center. Q4, quartile 4, are peaks with the strongest ChIP-seq peak signal, while q1, quartile 1, are peaks with the weakest ChIP-seq peak signal. Blue points indicate mA called with probability > 0.9, while orange points indicate mCpG called with probability > 0.9. Aggregate curves for each quartile were created with a 50 bp rolling window. Base density across the 2 kb region for each quartile is indicated. **b**, The distribution of differences between our single-molecule predicted peak center and the known CTCF motif are plotted for single molecules where we detected a peak. The distribution is fit to a Student’s t-distribution with 0.86 degrees of freedom. **c**, Joint mA and mCpG calls on the same individual molecules spanning the upper quartile of ChIP-seq peaks are displayed. Molecules displayed have at least one mA called and one mCpG called with probability > 0.9. Aggregate curves were created with a 50 bp rolling window. Base density is indicated. **d**, CTCF site protein occupancy is measured on single molecules spanning two neighboring CTCF motifs within 2-10 kb of one another. CTCF motifs are selected from quartile 3 and 4 ChIP-seq peaks, and molecules are shown that have a peak at at least one of the two motifs. Each row is a single molecule, and the molecules are anchored on the peaks that they span, with a variable distance between the peaks indicated by the grey block. ChIP-seq peak signal for each of the motif sites is indicated with the purple bars.

We next probed the relationship between CTCF binding and endogenous CpG methylation. Single molecules spanning CTCF binding sites in stronger ChIP-seq peaks exhibited a stronger dip in mCpG around the motif compared to the more shallow dip in weaker ChIP-seq peaks (Fig. 4a). This inverse relationship between CpG methylation and CTCF-targeted methylation as well as ChIP-seq strength reflects previous findings that mCpG inhibits CTCF binding (Bell and Felsenfeld 2000). Additionally, DiMeLo-seq provides sensitivity to detect this dynamic between mA and mCpG on single molecules (Fig. 4c). The in-phase periodicity of mA and mCpG reflects preferential methylation of both A and CpG within linker DNA. The increased methylation of CpG in linker DNA relative to nucleosome-bound DNA surrounding CTCF sites is supported by previous studies that have similarly reported higher levels of mCpG in linker DNA than nucleosomal DNA around CTCF sites (Kelly et al. 2012).

CTCF’s known specific binding motif and abundance genome-wide make it a good target for characterizing the resolution of DiMeLo-seq and for demonstrating the capability of DiMeLo-seq to detect joint binding events on single molecules. To characterize resolution, we called peaks on single molecules spanning the upper quartile of CTCF ChIP-seq peaks (Methods). We were able to predict the CTCF binding motif center within ~ 200 bp (−207 to 192 bp) on 70% of single molecules, and we found that the peak distribution center was shifted 8 bp 5’ from the motif center (Fig. 4b). This systematic bias towards predicting the peak center 5’ of the motif can be explained by the observed asymmetry in methylation when targeting CTCF. In addition to the accurate prediction of binding peak centers, another factor that impacts the resolution of DiMeLo-seq is the reach of the methyltransferase, which can be characterized by measuring the rate of methylation density decay from the peak center. To do this, we fit the adenine methylation density as a function of distance from the motif center to an exponential decay function and calculated a half-life of 125 bp (Supplementary Fig. 4b). To characterize the single-molecule sensitivity of our assay for detecting CTCF binding events, we calculated the fraction of reads that had at least one mA called with probability > 0.6 within 100 bp on either side of the motif center (Supplementary Fig. 4c). Using reads spanning the top decile of ChIP-seq peaks, we calculated DiMeLo-seq’s single-molecule sensitivity to be 40% (5% FDR, Methods).

We next investigated the ability of DiMeLo-seq to measure dynamics of protein binding at adjacent sites by characterizing CTCF occupancy across two binding sites spanned by a single molecule. We were able to detect neighboring CTCF motifs that are bound by CTCF at both sites, at one site, or at neither site (Fig. 4d). While the lack of signal at a site may be the result of the sensitivity of our assay rather than the vacancy of a CTCF site, this analysis demonstrates the potential to analyze coordinated binding patterns on single molecules. Unlike methods that detect protein binding across many molecules in aggregate, DiMeLo-seq can capture heterogeneity in joint occupancy across multiple binding sites from the same single molecule.

### 4. Mapping protein-DNA interactions in centromeric regions

#### Mapping histone modifications in heterochromatin with DiMeLo-seq

To test DiMeLo-seq’s ability to measure protein occupancy in heterochromatic, repetitive regions of the genome we targeted H3K9me3, which is abundant in pericentric heterochromatin. We chose to target H3K9me3 in HG002 cells because the chromosome X centromere has been completely assembled for this male-derived lymphoblast line (Nurk et al. 2021), and many different sequencing data types are available for it (Gershman et al. 2021). To validate the specificity of targeted methylation, we calculated the fraction of adenines methylated within HG002 CUT&RUN H3K9me3 peaks (Altemose et al. 2021) compared to the fraction of adenines methylated outside of broadly defined peaks (Methods). For H3K9me3 targeting in HG002 cells, the enrichment of mA/A in CUT&RUN peaks was 5.2-fold over background (Fig. 5a), indicating enrichment of methylation within expected H3K9me3-containing regions of the genome.

**Figure 5.**
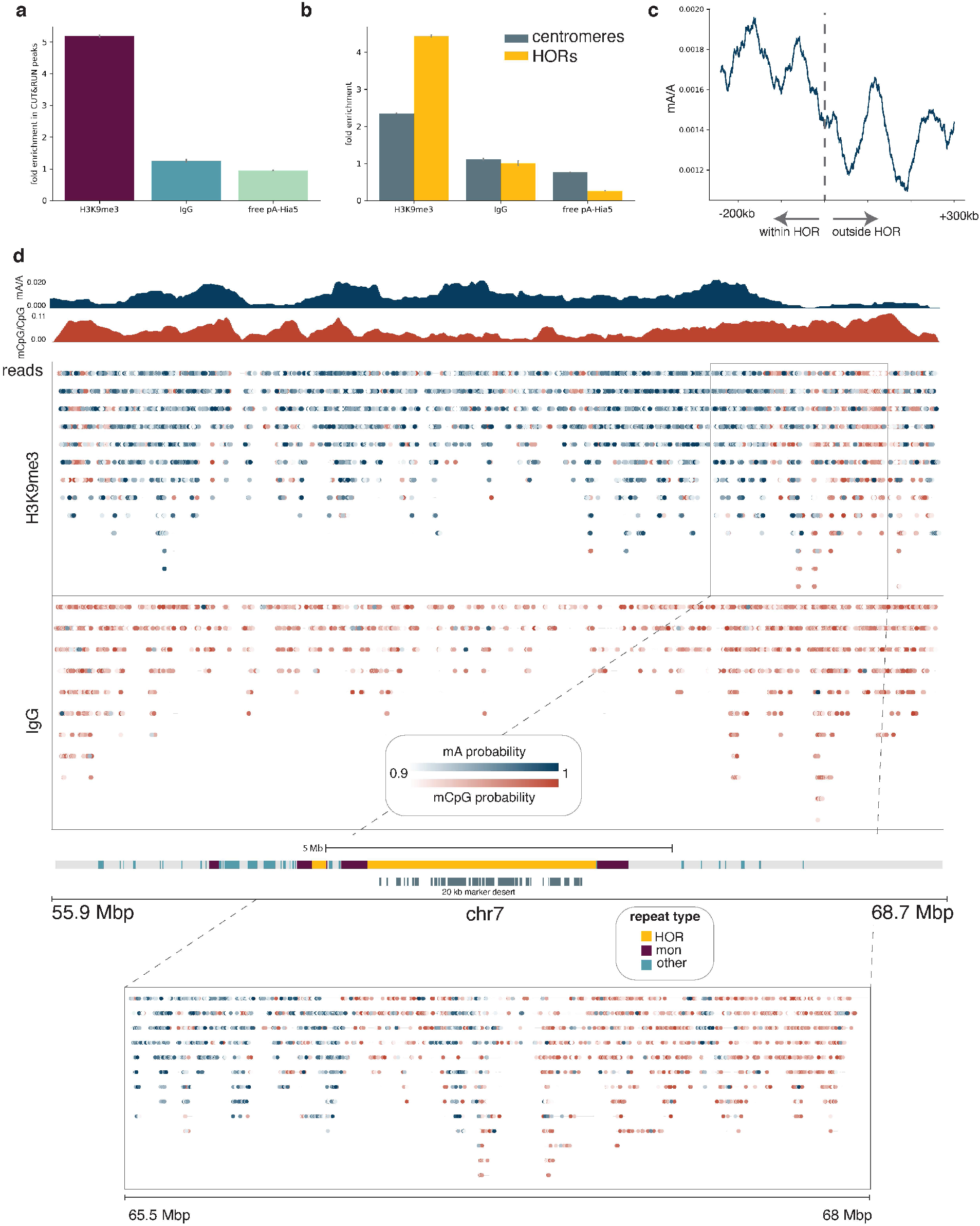
Detecting H3K9me3 in centromeres. **a**, The fraction of adenines methylated within CUT&RUN peaks relative to the fraction of adenines methylated outside of CUT&RUN broad peak regions is reported for the H3K9me3-targeted sample as well as IgG and free pA-Hia5 controls. Error bars represent the 95% confidence interval determined by simulating the proportion of adenines methylated as a Beta distribution with a uniform prior. **b**, The fraction of adenines methylated within centromeres relative to non-centromeric regions, and similarly the fraction of adenines methylated within active HOR arrays relative to non-centromeric regions are displayed for the H3K9me3-targeted sample as well as the IgG and free pA-Hia5 controls. Error bars are defined as in (a). **c**, The decline in mA/A for the H3K9me3-targeted sample in a rolling 60 kb window from −300 kb within the HOR array to 300 kb outside of the HOR array. HOR array boundaries that transition quickly into non-repetitive sequences were considered: 1p, 2pq, 6p, 9p, 13q, 14q, 15q, 16p, 17pq, 18pq, 20p, 21q, 22q. **d**, Single molecules are displayed across the centromere of chromosome 7 for the H3K9me3-targeted sample and the IgG control. Reads mapping to the same position are displayed vertically, and modified bases are colored by the probability of methylation at that base for probabilities > 0.9. Aggregate tracks show mA/A in the H3K9me3-targeted sample and mCpG/CpG across the H3K9me3-targeted sample and IgG and free pA-Hia5 controls in 10 kb bins. Grey bars below centromere annotation indicate regions with >20 kb marker deserts.

Human centromeres are located within highly repetitive alpha-satellite sequences, which are organized into higher order repeats (HORs) (McNulty and Sullivan 2018; Rudd, Schueler, and Willard 2003; Willard and Waye 1987; Altemose et al. 2021). To validate enrichment of H3K9me3-directed mA signal in centromeres, and in particular in HOR arrays, we similarly calculated the fold increase in mA/A and found 2.4-fold enrichment in centromeres and 4.4-fold enrichment in active (kinetochore-binding) HOR arrays (Altemose et al. 2021) over non-centromeric regions (Fig. 5b). We next looked at HOR array boundaries and observed a decrease in H3K9me3 across the boundary moving from within to outside of HOR arrays (Fig. 5c). In contrast, for the free pA-Hia5 control, mA/A increases moving from within to outside of the HOR array, as chromatin becomes more accessible (Supplementary Fig. 6a) (Gershman et al. 2021).

We mapped heterochromatin not only in aggregate across HOR array boundaries, but also in single molecules across the centromere. H3K9me3-targeted DiMeLo-seq reads map across the centromere of chromosome 7, even in regions with over 20 kb between unique markers (Fig. 5d). An IgG isotype control confirmed that adenine methylation in the H3K9me3-targeted sample was not caused by background methylation. Unlike methods which rely on amplifying short DNA fragments, such as ChIP-seq and CUT&RUN, we are able to detect single-molecule heterogeneity in chromatin boundaries, as highlighted in the transition from 65.5 Mb to 68 Mbp, where H3K9me3 signal drops as CpG methylation increases (Fig. 5d). However, low methylation efficiency in heterochromatin and limited mappability in repetitive regions can still lead to uneven and low coverage in these regions (Supplementary Fig. 6c). To improve sensitivity for targeted DiMeLo-seq applications in the centromere, we developed a centromere enrichment method to enhance coverage in active HOR arrays and applied this method to study CENP-A.

#### Restriction-based enrichment strategy improves centromere coverage

Within alpha satellite HOR arrays, the centromere-specific histone variant CENP-A delineates the site where the functional centromere and kinetochore will form. Population-level studies demonstrate that CENP-A nucleosomes are found at the core of these repeat units where the repeats are the most homogeneous (K. H. Miga et al. 2014; Hayden et al. 2013; Logsdon et al. 2021; Altemose et al. 2021). However, it has not been possible to resolve the positions of CENP-A nucleosomes on single chromatin fibers to determine the one-dimensional organization and density of CENP-A at centromeres. To map the positions of CENP-A nucleosomes at centromeres using DiMeLo-seq we developed a strategy to enrich specifically for human centromeric DNA in order to avoid sequencing the entire genome.

Our enrichment strategy, called AlphaHOR-RES (alpha higher-order repeat restriction and enrichment by size; from *alfajores*), is based on classic centromere enrichment strategies (Lica and Hamkalo 1983) that involve digesting the genome with restriction enzymes that cut frequently outside centromeric regions but rarely inside them, then removing short DNA fragments. Utilizing the gapless T2T-CHM13 human genome reference sequence (Nurk et al. 2021) to simulate digestion with a library of commercially available restriction enzymes, we identified two enzymes that, when combined, were predicted to enrich centromeric sequences over 20-fold among digestion products larger than 20 kb (MscI + AseI; Supplementary Fig. 7a). We added AlphaHOR-RES to our DiMeLo-seq workflow and observed at least 20-fold enrichment of sequencing coverage at centromeres while preserving relatively long read lengths (mean ~8 kb; Fig. 6a,b, Supplementary Fig. 7b-d, Methods). Thus, this enrichment strategy significantly increases the proportion of molecules sequenced that are useful for investigating CENP-A distribution, saving substantial sequencing time and costs. Furthermore, because AlphaHOR-RES targets the DNA and not the protein in the protein-DNA interaction, and because it is performed after directed methylation is complete, it is unlikely to bias our inferences of protein-DNA interaction frequencies in these regions.

**Figure 6:**
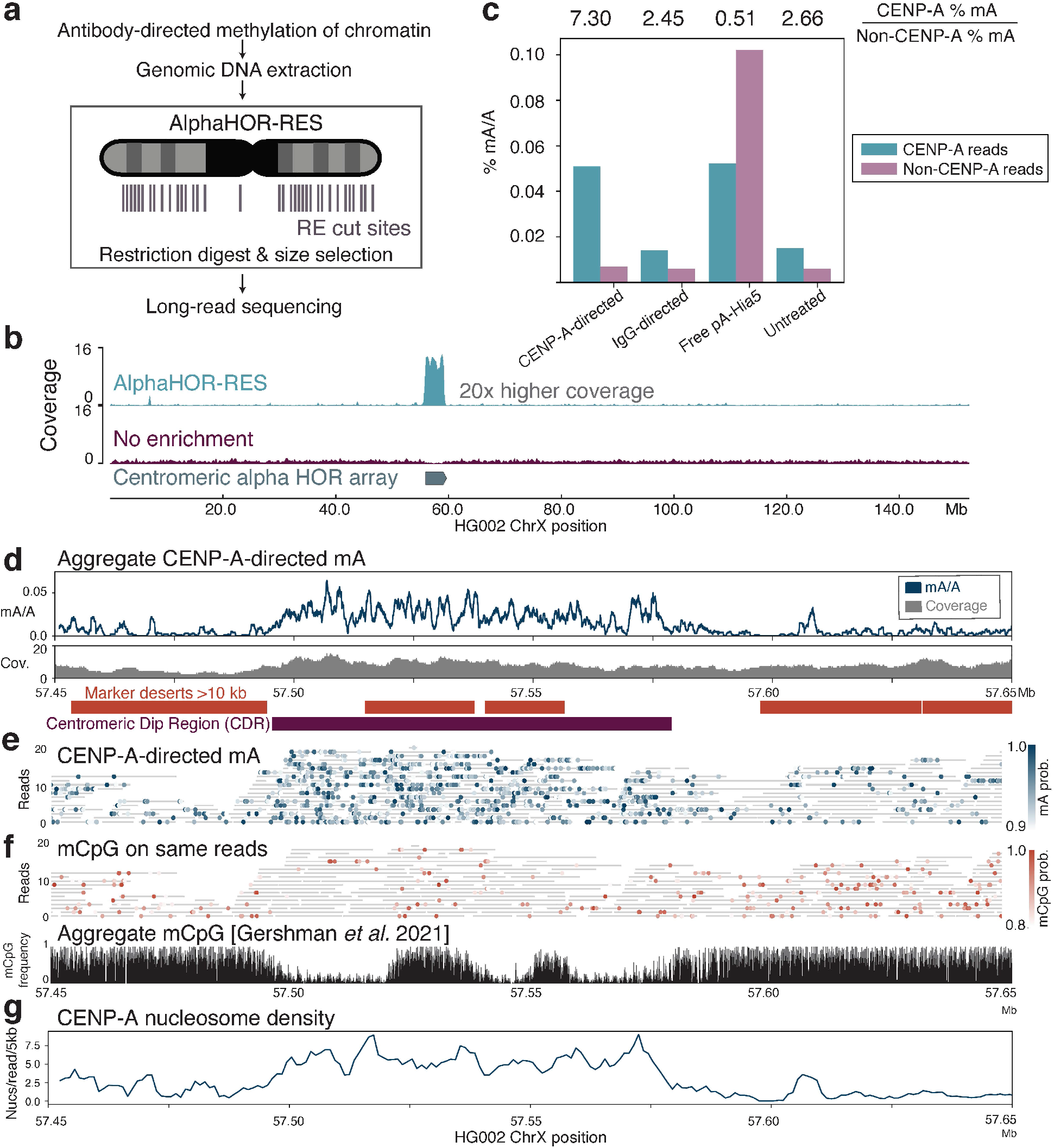
CENP-A directed methylation within chromosome X centromeric higher order repeats. **a**, Schematic of DiMeLo-seq with AlphaHOR-RES centromere enrichment. **b**, Genome browser plot on HG002 chromosome X of read coverage from DiMeLo-seq libraries with centromere enrichment (top) or without (middle). Bottom track depicts the region of alpha-satellite array. **c**, Barplot of percentage mA/A for reads from each library that contain (CENP-A reads) or do not contain (Non-CENP-A reads) CENP-A enriched *k*-mers. Fold enrichment of methylation percentage on CENP-A reads over non-CENP-A reads reported on top. **d**, Aggregate view of mA (top) and coverage (bottom) within a 200 kb region of chromosome X HOR spanning the CDR. mA/A plot indicates fraction of reads with methylation above threshold (0.9) probability (average over 1 kb rolling window for visualization). Regions of at least 10 kb without unique 51 bp *k*-mers are shown in orange to illustrate difficult-to-map locations for short-read sequencing. **e**, Single-molecule view with individual reads in gray and mA depicted as dots for CENP-A targeted DiMeLo-seq libraries. Scale bar indicates the probability of adenine methylation (from Guppy) between 0.9 and 1. **f**, (top) Single-molecule view of reads in **e**. showing endogenous mCpG depicted as dots. Scale bar indicates the probability of CpG methylation (from Guppy) between 0.8 and 1. (bottom) CpG methylation frequency from nanopore sequencing reported in (Gershman et al. 2021). **g**, Average number of nucleosomes estimated per read in sliding 5 kb windows for the portion of reads that align within the window (from reads that align to at least 1 kb within the window), to provide a measure of the density of CENP-A nucleosomes within single DNA molecules across the region.

#### DiMeLo-seq reveals variable CENP-A nucleosome density across centromeres

We performed CENP-A directed DiMeLo-seq on unfixed HG002 cells. After extraction of total genomic DNA, we used AlphaHOR-RES to enrich centromeric sequences before sequencing (Fig. 6a). In an alignment-independent manner, using sequence features enriched in CENP-A-targeted short-read sequencing datasets (Smith et al. 2021), we classified DiMeLo-seq reads based on the presence or absence of CENP-A-enriched *k*-mers. CENP-A-directed DiMeLo-seq reads with CENP-A enriched *k*-mers had ~7 fold higher percentage adenine methylation when compared to reads without CENP-A-enriched *k*-mers (Fig. 6c). We observed similar absolute methylation levels in DiMeLo-seq reads containing CENP-A *k*-mers when comparing CENP-A-targeted samples to free pA-Hia5 samples. However, the free pA-Hia5 samples also had a higher percentage of mA/A at reads that did not contain CENP-A *k*-mers, indicating a lack of CENP-A specificity in the absence of targeting.

To examine the positions of CENP-A nucleosomes within centromeric repeat arrays, we aligned our reads to a hybrid complete human assembly containing a fully assembled chromosome X from HG002 (Methods) (Nurk et al. 2021; Gershman et al. 2021). We investigated the recently described chromosome X centromere dip region (CDR), a hypomethylated region in the centromeric alpha HOR array where short-read CENP-A datasets align (Gershman et al. 2021; Logsdon et al. 2021; Karen H. Miga et al. 2020; Altemose et al. 2021). Aggregating the methylation signal across multiple reads, we confirmed low endogenous CpG methylation within that region as expected (Fig. 6f). CENP-A-directed mA was found to be higher within both large and small CDRs compared to their adjacent CpG methylated regions, consistent with short-read data for this cell line (Fig. 6d-f) (Altemose et al. 2021; Gershman et al. 2021). Furthermore, we found that the number of CENP-A nucleosomes detected per kilobase on single-molecule reads increased over 4-fold within ChrX CDRs compared to neighboring regions (Methods, Fig. 6e,g), confirming what ensemble short-read methods cannot: the *density* of CENP-A nucleosomes on single DNA molecules increases in CDRs. It is important to note, however, that an absolute quantification of CENP-A nucleosome density in centromeres will require further characterization of the single-molecule sensitivity of DiMeLo-seq in these regions, which will provide a new way to test previous estimates made by orthogonal methods (Bodor et al. 2014). IgG isotype controls confirm that this signal is not due to background methylation (Supplementary Fig. 8a). While CENP-A-directed methylation is enriched in the CDR as a whole, individual molecules have distinct methylation patterns that do not appear stereotyped, suggesting dense but potentially variable placement of CENP-A nucleosomes within this region. We observe a similar distribution of CENP-A-directed methylation on chromosome 3, where only one of the two HOR arrays was observed to have clear CENP-A-directed methylation (Supplementary Fig. 8b,c). These observations support the finding of one active HOR array per chromosome (Aldrup-MacDonald et al. 2016; Altemose et al. 2021). We also observe single-cell heterogeneity in the distribution of CENP-A nucleosomes, and their relation to CpG methylation, noting variegation in transitions between regions with high and low CENP-A density (Fig. 6 e-f). These findings illuminate the density and positioning of CENP-A nucleosomes within HOR sequences on individual chromatin fibers, which was not previously attainable with existing techniques.

## Discussion

Here, we have developed, optimized, and validated DiMeLo-seq, a new method for mapping protein-DNA interactions genome-wide. DiMeLo-seq can map a protein’s binding sites within hundreds of base pairs at multiple loci on single molecules of sequenced DNA up to hundreds of kilobases in length. This long read length improves mappability in repetitive regions of the genome, opening repeat regions up to new studies of their regulation. Because DiMeLo-seq involves no amplification, it can provide a proportional readout of protein-DNA binding frequency at every site. It can also provide joint information about endogenous CpG methylation and protein-DNA interactions on the same long single molecules. We produced an optimized DiMeLo-seq protocol that only requires ordinary lab equipment to prepare libraries (Supplementary Fig. 1). This protocol is also compatible with cryogenically frozen and lightly fixed samples, expanding the range of potential samples and targets (Supplementary Table 1; interactive, updated protocol on protocols.io; see Methods).

Using a controlled reconstituted chromatin system, we showed that DiMeLo-seq can direct methylation to specific histone protein targets and methylate adenines on the surrounding linker DNA, revealing the footprint of each targeted nucleosome on single DNA molecules. We mapped interactions between DNA and the nuclear lamin protein LMNB1 to evaluate DiMeLo-seq’s signal-to-background ratio (up to 30-fold), its single-molecule sensitivity (up to ~50%), and its ability to proportionally detect known interaction frequencies. We also mapped CTCF, a zinc-finger protein with a small binding footprint, and showed that (i) we can resolve its binding sites within hundreds of base-pairs on single molecules, (ii) we can observe reduced CpG methylation at its binding sites on the same single molecules, and (iii) we can jointly map multiple binding events on single molecules. We demonstrated our ability to methylate and map protein-DNA interactions within highly repetitive centromeric regions by targeting both H3K9me3 and CENP-A, and we created alphaHOR-RES, a method for enriching centromeric DNA 20-fold to improve coverage and cost-effectiveness. Finally, by mapping individual CENP-A nucleosomes on long, sequenced DNA molecules for the first time, we found that: (i) CENP-A nucleosome density increases on single chromatin fibers in mCpG depleted regions within centromeres, and (ii) CENP-A nucleosomes lack evidence of having highly stereotyped positioning. Because pre-existing CENP-A nucleosomes are thought to epigenetically direct the assembly of new CENP-A nucleosomes in each cell cycle, it will be interesting to understand how this seemingly stochastic distribution arises along the sequence of the active centromere.

This study also allowed us to characterize the benefits and tradeoffs of using DiMeLo-seq compared to short-read ensemble methods. Because DiMeLo-seq is an amplification-free method that sequences single native DNA molecules, and because it relies on centrifugation for washing steps, it requires a relatively large amount of starting material (1-2 million cells per replicate). Using concanavalin-A coated magnetic beads, which we demonstrated to be compatible with the DiMeLo-seq protocol, may help to reduce these input requirements (as in certain variations of CUT&RUN) (Skene and Henikoff 2017). Within a particular bin of the genome, each read can be thought of as a single-cell data point, but different bins across the genome cannot be jointly analyzed in the same cell because single-cell coverage is extremely low without amplification. Additionally, the lack of enrichment means that the standard DiMeLo-seq protocol requires the entire genome to be sequenced uniformly, potentially wasting sequencing reads in regions of the genome that are irrelevant for the target protein’s binding domain. For proteins that only target small regions, it is possible to perform targeted DNA sequencing (Gilpatrick et al. 2020; Kovaka et al. 2021) or to use DNA enrichment methods like alphaHOR-RES, the centromere enrichment method we demonstrated here. It is also possible to use immunoprecipitation to enrich for methylated DNA or DNA bound to a protein of interest, but this would destroy the proportionality between methylation frequency and protein-DNA interaction frequency. We estimated the single-molecule sensitivity of DiMeLo-seq to be between 40-50% (for CTCF and LMNB1), though this may vary by target protein and antibody, perhaps owing to inefficiencies in cell permeabilization and methyltransferase activity, or to single-molecule variability in local steric effects or in the binding strength of the target protein, antibody, or pA. We expect this sensitivity can be improved by further optimization and extension of this method. However, because DiMeLo-seq samples DNA molecules proportionally, once the sensitivity for a particular target has been estimated, this estimate can be used to adjust inferences of phenomena like joint binding frequencies on single molecules.

Because Hia5 only tends to methylate unbound linker DNA, DiMeLo-seq provides information about local nucleosome occupancy along with the target protein’s footprint. This also means that highly inaccessible regions can be more difficult to methylate, and they may require higher sequencing coverage. Additionally, because DiMeLo-seq is performed *in situ* in conditions meant to preserve chromatin conformation, it may methylate unbound DNA in *trans* if it is close enough to the target protein’s binding sites in 3D space, as does CUT&RUN (Skene and Henikoff 2017). These 3D interactions, and the factors that mediate them, can potentially be investigated by perturbing 3D chromatin structure prior to performing DiMeLo-seq, which may also be a useful approach for improving DNA accessibility in highly condensed regions.

We anticipate that DiMeLo-seq will be useful for investigating a wide range of biological questions. For example, because it can allow one to explore the density of a protein’s binding along a single chromatin fiber from a single cell, it can be used to investigate how the exact boundaries between chromatin states vary among single cells, or perhaps how the stoichiometry of a DNA-binding protein in enhancers affects the transcription of nearby genes. The method is also compatible with *in vivo* expression of protein-MTase fusions, as in conventional DamID (van Steensel and Henikoff 2000), instead of antibody targeting *in situ*. This would prove useful for investigating more transient protein-DNA interactions, or proteins that lack suitable antibodies. One can also imagine adding exogenous GpC methylation marks to provide information about a second protein’s joint binding profile. Another potential advantage is that because protein-DNA interactions are measured simultaneously with sequence on long reads, DiMeLo-seq provides the opportunity to map protein-DNA interactions while performing *de novo* genome assembly with no additional sequencing cost, or to investigate the relationship between sequence variation and protein binding within phased single molecules. As nanopore sequencing and other long-read sequencing platforms become more efficient, more accurate, and cheaper, DiMeLo-seq will extend these technologies for more comprehensive mapping of protein-DNA interaction landscapes.

## Acknowledgements

We thank Andrew Stergachis for the plasmid encoding Hia5 and Gary Karpen for helpful discussions. We would like to thank Stanford University and the Stanford Research Computing Center for providing computational resources and support that contributed to these research results. This work was supported by the Chan Zuckerberg Biohub and by the NIGMS of the National Institutes of Health under award number R35GM124916 to AS and R01 GM074728 to AFS. OKS and RRB are supported by an NIH T32 award, numbers GM113854-02 and GM007279-45 respectively. AM, OKS, and RRB are supported by NSF GRFP awards. NA is an HHMI Hanna Gray Fellow. AS is a Chan Zuckerberg Biohub Investigator, and a Pew Scholar in the Biomedical Sciences.

## Author Contributions

NA, AM, OKS, KS, AFS, and AS designed the study. NA, AM, OKS, KS, and RRB performed the experiments. AD and NN assisted with sequencing. KHM provided unpublished datasets and feedback. NA, AM, OKS, and KS analyzed and interpreted the data. NA, AM, OKS, KS, and RRB made the figures. NA, AM, OKS, and KS wrote the manuscript, with input from RRB, AFS, and AS. AFS and AS supervised the study.

## Methods

### Protocols/Data/Code/Materials availability

For detailed and updated protocols, please refer to the following protocols.io web pages:

**DiMeLo-seq**: https://dx.doi.org/10.17504/protocols.io.bv8tn9wn

**pA-Hia5 Protein Purification**: https://dx.doi.org/10.17504/protocols.io.bv82n9ye

**AlphaHOR-RES**: https://dx.doi.org/10.17504/protocols.io.bv9vn966

All sequencing data are available by request and will be made available on GEO upon publication.

All data analysis code is available by request and will be available on github upon publication.

pA-Hia5 expression plasmid ASP4201 and pAG-Hia5 expression plasmid ASP4221 are available by request, and they will be deposited to Addgene prior to publication.

### Cell culture

HEK293T cells (CRL-3216, ATCC, Manassas, VA; validated by microsatellite typing and mycoplasma tested) were maintained in DMEM (high glucose, with GlutaMAX, with phenol red, without sodium pyruvate; Gibco 10566016) supplemented with 10% Fetal Bovine Serum (VWR 89510-186) and 1% Pen Strep (Gibco 15070063) at 37°C in 5% CO_2_. GM12878 cells (Coriell Institute, Camden, NJ; mycoplasma tested) and HG002 cells (Coriell Institute, Camden, NJ; mycoplasma tested) were maintained in RPMI-1640 with L-glutamine (Gibco 11875093) supplemented with 15% Fetal Bovine Serum (VWR 89510-186) and 1% Pen Strep (Gibco 15070063) at 37°C in 5% CO_2_.

### Cloning of pA-pHia5ET and pAG-pHia5ET

The pHia5ET vector was generously provided by Andrew Stergachis and John Stamatoyannopoulos (Stergachis et al. 2020). Protein A (pA) was amplified from pK19pA-MN (ASP4062, Addgene plasmid #86973, ref: (Schmid, Durussel, and Laemmli 2004)) and Protein AG (pAG) was amplified from pAG/MNase (ASP4154, Addgene plasmid #123461, ref: (Meers et al. 2019)). The pHia5ET vector was linearized via NdeI restriction digest. pA or pAG was inserted in front of the Hia5 cassette in pHia5ET using Gibson Assembly. All plasmid sequences were verified using Sanger sequencing.

### Protein purification

Histones for chromatin assembly (CENP-A, H3, H4, H2A, and H2B) were purified as previously described ((Guse, Fuller, and Straight 2012), (Westhorpe, Fuller, and Straight 2015)). pA-Hia5, pAG-Hia5, and Hia5 purification were adapted from (Stergachis et al. 2020). Plasmids were transformed into T7 Express lysY competent *E. coli* cells (NEB #C3010I) for recombinant protein expression. 200 mL starter culture was grown in LB broth at 37°C with 50 ug/mL kanamycin and 34 ug/mL chloramphenicol to an OD_600_ of 0.6. Starter cultures were then diluted to 2L culture in LB broth at 37°C with 50 ug/mL kanamycin to an OD_600_ of 0.8 - 1.0. Protein expression was induced using a final concentration of 1mM IPTG (Isopropyl beta-D-1-thiogalactopyranoside) for 4 hours at 20°C with shaking. Cells were then pelleted at 5000 x g for 15 minutes at 4°C. Pelleted cells were resuspended in 35 mL lysis buffer (50 mM HEPES, pH 7.5; 300 mM NaCl; 10% glycerol; 0.5% Triton X-100). Resuspended cell pellets were flash frozen in liquid nitrogen and stored at −80°C until purification. After thawing frozen cell pellets, EDTA-free protease inhibitor tablets (Roche 11873580001) and 10 mM β-mercaptoethanol were added. Cells were lysed by probe sonication (6 pulses, 30s on, 1 min off at 200W). Lysed cells were centrifuged for 1 hour at 40,000 x g (4°C) in 50 mL Oakridge tubes. Ni-NTA agarose was prepared with 2x washes of 30 mL equilibration buffer (50 mM HEPES, pH 7.5; 300 mM NaCl; 20 mM imidazole) per 5 mL of slurry. Cell lysate was incubated with Ni-NTA agarose and rotated for 1 hour at 4°C. Mixture was poured onto a gravity column, then washed with 40 mL buffer 1 (50 mM HEPES, pH 7.5; 300 mM NaCl; 50 mM imidazole), 30 mL of buffer 2 (50 mM HEPES, pH 7.5; 300 mM NaCl; 70 mM imidazole), and eluted with 30 mL of elution buffer (50 mM HEPES, pH 7.5; 300 mM NaCl; 250 mM imidazole). Eluted protein was filtered with a 0.2 μm filter, then loaded onto a HiPrep 26/10 Desalting column (Cytiva) to buffer exchange eluate into FPLC buffer A (50 mM Tris-HCl, pH 8.0; 100 mM NaCl; 1mM DTT). Following buffer exchange, the sample was applied in tandem onto HiTrap Q HP and HiTrap SP HP (Cytiva) columns. Both columns were washed with 5x combined column volumes of FPLC buffer A. The HiTrap Q HP (Cytiva) column was removed and protein was eluted from the SP column using a linear gradient of 20 column volumes with increasing linear gradient of FPLC buffer B (50 mM Tris-HCl, pH 8.0; 1 M NaCl; 1 mM DTT). Fractions were collected and quantified using A280 absorbance. Elution peak fractions were concentrated using a 10K Amicon Ultra-15 tube to final protein concentration > 5 uM. The final concentrated protein was supplemented with 10% glycerol final concentration, aliquoted, and stored at −80°C.

### DiMeLo-seq

All reagents were prepared fresh, syringe filtered through a 0.2 μm filter, and kept on ice. Cells (1M-5M per condition) were pelleted at 300 × g for 5 minutes and washed with PBS. While live cells were used for experiments targeting CTCF, H3K9me3, CENP-A, and the accompanying controls, both frozen and fixed cells are also compatible with the DiMeLo-seq protocol. Frozen cells stored in freezing medium with DMSO in liquid nitrogen should be thawed on ice and prepared with the same protocol as fresh cells. For optional light fixation, cells can be fixed with 0.1% PFA for 2 minutes with gentle vortexing, followed by the addition of 1.25 M glycine to twice the molar concentration of PFA, a 3 minute spin at 500 x g at 4°C, and then continuation with standard DiMeLo-seq protocol’s nuclear isolation. Pelleted cells were resuspended in 1 ml of Dig-Wash buffer (0.02% digitonin, 20 mM HEPES-KOH, pH 7.5, 150 mM NaCl, 0.5 mM Spermidine, 1 Roche Complete tablet -EDTA (11873580001) per 50 ml buffer, 0.1% BSA) and incubated on ice for 5 minutes. The nuclei suspension was then split into separate tubes for each condition and spun down at 4°C at 500 x g for 3 minutes. All subsequent spins were performed with these same conditions, and all steps involving pipetting nuclei were performed with wide bore tips. The supernatant was removed and the pellet was gently resolved in 200 μl Tween-Wash (0.1% Tween-20, 20 mM HEPES-KOH, pH 7.5, 150 mM NaCl, 0.5 mM Spermidine, 1 Roche Complete tablet-EDTA per 50 ml buffer, 0.1% BSA) containing the primary antibody at a 1:50 dilution. Antibodies targeted the following: LMNB1 (ab16048), CTCF (ab188408), H3K9me3 (Active Motif 39162), CENP-A (Aaron Straight, Stanford University, (Carroll et al. 2009)), and rabbit IgG isotype control (ab171870). Samples were placed on a rotator at 4°C for 2 hours. Nuclei were then pelleted and washed twice with 0.95 ml Tween-Wash. For each wash, the pellet was completely resolved by pipetting up and down ~10 times and placed on a rotator at 4°C for 5 minutes before spinning down. Following the second wash, the nuclei pellet was gently resolved in 200 μl Tween-Wash containing 200 nM pA-Hia5. pA-Hia5 concentration was measured using the Qubit Protein Assay Kit (Q33211). For pA-Hia5 binding, the nuclei were placed on the rotator at room temperature for 1 hour. Nuclei were then spun down and washed twice with 0.95 ml Tween-Wash with a 4°C rotating incubation for 5 minutes between spins, as in the wash following antibody binding. For the free pA-Hia5 control, nuclei were kept on the rotator at 4°C during antibody binding and pA-Hia5 binding steps, and pA-Hia5 was added at the time of activation. Nuclei were then resuspended in 100 μl of Activation Buffer (15 mM Tris, pH 8.0, 15 mM NaCl, 60 mM KCl, 1 mM EDTA, pH 8.0, 0.5 mM EGTA, pH 8.0, 0.5 mM Spermidine, 0.1% BSA, 800 μM SAM) and incubated at 37°C for 30 minutes before spinning and resuspending in 100 μl of cold PBS. Depending on the desired read length, either the Monarch Genomic DNA Purification Kit (T3010S) or the NEB Monarch HMW DNA Extraction Kit (T3050L) with 2000 rpm agitation was used to extract DNA from the nuclei. If fixation was performed, incubate at 56°C for 1 hour for lysis to reverse crosslinks. For T3050L, agitate for the first 10 minutes of lysis and then keep the samples at 56°C without agitation for 50 minutes. DNA yield was quantified using the Qubit dsDNA BR Assay Kit (Q32850).

Immunofluorescence imaging following binding with pA/G-MTase (i.e. pA-Hia5 or pAG-Hia5 or pAG-EcoGII) was performed to evaluate cell permeabilization, nuclear integrity, primary antibody on-target and background binding. For detection of pA/G-MTase binding, two different fluorophore-conjugated antibodies were used: a goat anti-mouse IgG antibody conjugated to AlexaFluor647 (Invitrogen A32728), which is not expected to bind to the rabbit primary or goat secondary antibodies but is expected to be bound by pA/G, and a goat anti-V5 antibody conjugated to FITC (Abcam 1274), which is expected to bind to the C-terminal V5 tag on pA/G-MTase. It is also possible to use a chicken anti-HisTag FITC-conjugated antibody (Abcam 3554) to avoid any binding by pA or pG.

### Nanopore library preparation and sequencing

For each sample, 3μg DNA was input into library preparation using one of the following library preparation kits: (1) Ligation Sequencing Kit (ON SQK-LSK109) with Native Barcoding Expansion 1-12 (ON EXP-NBD104) and Native Barcoding Expansion 13-24 (ON EXP-NBD114) for optimization experiments and CENP-A targeted experiments after AlphaHOR-RES, or (2) Ligation Sequencing Kit (ON SQK-LSK110) for CTCF targeting, H3K9me3 targeting, and the corresponding IgG and free pA-Hia5 controls in GM12878 and HG002.

For method (1), the protocol was performed as described in the LSK109 documentation with the following modifications. End repair incubation time was increased to 10 minutes. 1 μg of end repaired DNA was loaded into barcode ligation. All ligation incubation times were increased to at least 20 minutes. Elution following barcode ligation reaction cleanup was decreased to 18 μl to allow for loading 3 μg of pooled barcoded material into the final ligation. If DNA was not sufficiently concentrated, the speedvac was used to concentrate the DNA. LFB was used for the final cleanup and elution was performed with 13 μl EB. 1 μg of DNA was loaded onto the sequencer.

For method (2), initial runs following high molecular weight extraction using NEB Monarch HMW DNA Extraction Kit with 2000 rpm agitation during lysis suffered from bead clumping during library preparation cleanups, resulting in low yields and reduced fragment size. To preserve longer fragments with the LSK110 kit, the following modifications were made to the standard LSK110 protocol (Kim et al. 2020). End preparation incubation time was increased to 1 hour with a 30 minute deactivation. The cleanup following end preparation was performed by combining 60 μl SRE buffer (Circulomics SS-100-101-01) with the 60 μl end prep reaction, centrifuging at 10,000 x g at room temperature for 30 minutes, or until the DNA had pelleted, and washing with 150 μl 70% ethanol two times with a 2 minute spin at 10,000 x g between washes. The pellet was resuspended in 31 μl EB, and incubated at 50°C for 1 hour and then 4°C for at least 48 hours. Ligation volume was reduced by half for a total of 30 μl DNA in a 50 μl reaction volume. The ligation incubation was increased to 1 hour. The DNA was pelleted at 10,000 x g at room temperature for 30 minutes. The pellet was washed twice with 100 μl LFB, with a 2 minute spin at 10,000 x g between washes. The pellet was resuspended in 31 μl EB and incubated at least 48 hours at 4°C. For sequencing, 500 ng of the final library was loaded, with a wash using the Flow Cell Wash Kit (ON EXP-WSH004) and reload every 24 hours. Other approaches, such as using Zymo Genomic DNA Clean & Concentrator (D4065) for cleanup between reaction steps in the LSK110 protocol and and using the Rapid Barcoding Kit (ON SQK-RBK004) were performed; however, LSK110 with pelleting DNA for cleanup resulted in the best throughput with the longest reads.

Sequencing was performed on an Oxford Nanopore MinION sequencer with v9.4 flow cells (ON FLO-MIN106.1). N50 varied with library preparation method, with a range from ~30 kb with LSK110 without modification to ~50-70 kb with LSK110 with the modifications for pelleting for DNA cleanup. See Supplementary Table 3 for summary sequencing metrics for each sample.

### Reconstituted chromatin experiments

18×601 DNA array was obtained as previously described (Guse, Fuller, and Straight 2012). To summarize, puC18 vector with 18 repeats of the “601” nucleosome positioning sequence (Lowary and Widom 1998) (ASP 696) was transformed into competent dam-*E. coli* strain, GM2163, and purified using a QIAGEN Gigaprep kit. The unmethylated 18×601 plasmid was digested with EcoRI, XbaI, DraI, and HaeII. Array DNA used in directed methylation experiments was then biotinylated by filling in EcoRI and XbaI 5’overhangs with dGTP (NEB), α-thio-dCTP (Chemcyte), α-thio-dTTP (Chemcyte), and biotin-14-dATP (Thermo Fisher Scientific) using Large Klenow fragment 3’-5’ exo- (NEB).

Chromatin was reconstituted using salt dialysis as described in Guse et al 2012. 18×601 DNA, H2A/H2B histone dimer, and tetramer (H3/H4 or CENP-A/H4 histone tetramer) were added to high salt buffer (10 mM Tris-HCl, pH 7.5; 0.25 mM EDTA; 2 M NaCl). The mixture was gradually dialyzed over the course of ~67 hours at a rate of 0.5 mL/min from high salt buffer into low salt buffer (10 mM Tris-HCl, pH 7.5; 0.25 mM EDTA; 2.5 mM NaCl). CENP-A/H4 or H3/H4 tetramer concentrations were titrated to obtain chromatin of varying saturation. Chromatin assembly was verified using a native acrylamide gel shift analysis after overnight restriction digestion using AvaI at room temperature (18×601 array DNA contains engineered AvaI recognition sites between adjacent 601 positions)(Guse, Fuller, and Straight 2012).

### In vitro DNA methylation assay

Hia5, pA-pHia5, and pAG-pHia5 concentrations were estimated using the extinction coefficients. Serial dilutions (100, 10, 1, 0.1 nM for Hia5 and pA-Hia5 comparison, 30, 3, 0.3 nM for Hia5 and pAG-Hia5 comparison) were made using Buffer A (15 mM Tris-HCl, pH 8.0; 15 mM NaCl, 60 mM KCl, 1 mM EDTA, 0.5 mM EGTA, and 0.5 mM spermidine). Proteins were then mixed with Buffer A supplemented with S-adenosyl-methionine (NEB B9003S) containing 1 ug of either naked unmethylated DNA (Plasmid ASP3552, 2×601, prepared from GM2163 dam-*E. coli* strain) or methylated DNA (Plasmid ASP3552, 2×601, prepared from DH5a *E. coli* strain). Reactions were incubated for 1 hr at 37°C. PCR purification was performed to extract DNA which was then digested with DpnI for 1.5 hours at 37°C and run on an agarose gel to assess the degree of methylation.

### In vitro chromatin methylation

In experiments involving free pA-Hia5 (non-targeted) methylation on chromatin, reconstituted chromatin (at 12.5 nM 601 concentration or 0.7 pM 18×601 concentration) was incubated in activation buffer (15 mM Tris-Cl pH 8.0, 15 mM NaCl, 60 mM KCl, 0.1% w/v Bovine Serum Albumin (BSA)) containing 0.8 mM SAM and 25 nM pA-Hia5 or pAG-Hia5 for 30 minutes at 37 °C. In antibody-directed methylation experiments, chromatin reconstituted on biotinylated 18×601 array DNA was used. In DNA LoBind Eppendorf tubes, M-280 Streptavidin-coated Dynabeads (Invitrogen) were washed in bead buffer (50 mM Tris, pH 7.4, 75 mM NaCl, 0.25 mM EDTA, 0.05% Triton X-100, and 2.5% polyvinyl alcohol (30kDa - 70 kDa)) and incubated with biotinylated CENP-A or H3 containing chromatin (at 12.5 nM 601 concentration or 0.7 pM 18×601 concentration) for 1 hour at room temperature with constant agitation. Chromatin-coated beads were then magnetically separated and washed twice with Chromatin wash buffer (CWB)-75 (20 mM HEPES pH 7.5, 75 mM KCl, 0.05% Triton X-100, 0.1% BSA) and then incubated in CWB-75 containing 1 ug/mL of rabbit anti-CENP-A antibody (Carroll et al. 2009), mouse anti-H3 antibody (MABI 0301, Active Motif), or rabbit or mouse IgG (Invitrogen) control for 30 minutes with agitation at room temperature. Beads were then washed twice with CWB-75 and incubated in CWB-75 for 30 minutes with agitation, followed an additional wash in CWB-75 before incubation in CWB-75 containing 25 nM pA-Hia5 (in CENP-A-directed methylation experiments) or pAG-Hia5 (in H3-directed methylation experiments). After incubation with pA-Hia5 or pAG-Hia5, beads were washed twice with CWB-100 (20 mM HEPES pH 7.5, 100 mM KCl, 0.05% Triton X-100, 0.1% BSA) to remove unbound pA-Hia5 and then resuspended in activation buffer (15 mM Tris-Cl pH 8.0, 15 mM NaCl, 60 mM KCl, 0.1% w/v BSA) containing 0.8 mM SAM for

30 minutes at 37°C. Beads were then split into two tubes and processed separately for immunostaining (with anti-mA antibody) and library preparation (for long-read sequencing). For library preparation, chromatin was released from beads using BamHI and KpnI digestion (cuts near biotinylated ends of 18×601 array DNA), DNA was extracted, and processed using Oxford Nanopore Technology native barcoding (PCR-free) kit (EXP-NBD104 or EXP-NBD114) with the Ligation Sequencing Kit (SQK-LSK109).

### Chromatin-coated beads immunostaining and imaging

Following incubation in activation buffer, chromatin coated beads were incubated in CWB-2M (20 mM HEPES pH 7.5, 2M NaCl, 0.05% Triton X-100, 0.1% BSA) for 1 hour at 55 °C to denature protein. Beads were then washed twice in CWB-2M to remove denatured protein while retaining biotinylated DNA on beads. Beads were then washed twice with Antibody dilution buffer or AbDil (20 mM Tris-HCl, pH 7.4, 150 mM NaCl with 0.1% Triton X-100, and 2% BSA) and dropped onto poly-L-lysine-coated coverslips and allowed to attach for 30 minutes. Coverslips were incubated with AbDil containing 1 ug/ml rabbit anti-N6-methyladenosine antibody (Millipore Sigma ABE572) for 30 minutes, washed twice with AbDil, and incubated with AbDil containing 2 ug/ml Alexa 647 fluorophore conjugated goat anti-rabbit secondary antibody (Molecular Probes) for 30 minutes. Coverslips were then washed twice with AbDil, incubated with AbDil containing 1 ug/ml propidium iodide (Sigma) for 10 minutes, washed twice with AbDil and phosphate buffered saline (PBS), blotted gently, mounted in 90% glycerol, 10 mM Tris-Cl pH 8.8, and 0.5% *p*-phenylenediamine, and sealed using clear nail polish.

Imaging was performed using IX70 (Olympus) microscope with a DeltaVision core system (Applied precision) with a Sedat quad-pass filter set (Semrock) and monochromatic solid-state illuminators, controlled via softWoRx 4.1.0 software (Applied Precision). Images were acquired using a 100x 1.4 NA Plan Apochromatic oil immersion objective (Olympus) and captured using a CoolSnap HQ CCD camera (Photometrics). Z-stacks were acquired at 0.2 uM intervals over a total 3 uM total axial distance. Bead images were analyzed using custom ruby software (Westhorpe, Fuller, and Straight 2015). At least 50 beads were analyzed for each condition per experiment.

### In vitro chromatin DiMeLo-seq analyses

Reads from *in vitro* experiments were initially basecalled with Guppy (4.4.2) using the fast basecalling model (dna_r9.4.1_450bps_fast.cfg). After initial basecalling reads were demultiplexed and split by barcode using the guppy_barcoder and fast5_subset from ont_fast5_api. Fast5s for each barcode were then aligned and modification basecalled with Megalodon (2.2.9) using the rerio all-context basecalling model (res_dna_r941_min_modbases-all-context_v001.cfg) with --guppy_params “trim_barcodes” and --mod_min_prob 0. Modification basecalled reads were smoothed by calculating rolling average over a 33 bp window in a Nan-sensitive manner (averaging only over adenine bases). Following smoothing, adenine bases with methylation probability score > 0.6 were assigned as methylated (mA) (threshold of 0.6 was empirically determined by comparing pA-Hia5 treated and untreated naked 18×601 DNA, false positive rate = 3%). Clustering of reads were performed using hierarchical clustering of jaccard distances of mA positions within the 18×601 or 4×601 portion as described in figure. For measuring distance of gaps from the theoretical 601 dyad axis, continuous regions without mA (or gaps) of 100 - 180 bp lengths were filtered and the offset of the center of these gaps from the nearest 601 dyad position was calculated. For measurements of average mA probability score at and near gaps, 100 - 180 bp sized gaps that lack 100 - 180 bp sized gaps within 400 bp (approximately 2x 601 repeat lengths) on either side were extracted, centered by the gap centers and the average Megalodon probability score was calculated at each position.

### Centromere enrichment using alphaHOR-RES

The T2T-CHM13v1.0 reference genome was *in silico* digested with all 4-6 bp restriction enzymes available from New England Biolabs annotated as insensitive to dam or CpG methylation. A subset of these enzymes were selected based on the criteria of having less than 5% of the generated fragments map back to the alpha-satellite region of the genome and for which the genome was fragmented into at least 200,000 total fragments. Centromere enrichment was calculated after artificially removing fragments under 20 kb to simulate a size selection step and determining the fraction of remaining fragments that map to centromeric regions, as well as the loss of alpha satellite containing sequences (Supplementary Fig. 7). Combinations of digests were then evaluated and MscI and AseI were identified as an optimal pair for centromere enrichment.

Genomic DNA was extracted from ~25 million cells using an NEB HMW DNA extraction kit using 300 rpm rotation during lysis (#T3050L). The DNA was eluted in a total of 300 μl elution buffer and allowed to relax at 4 °C for 2 days, although it remained viscous until it was solubilized. 37 μl NEBuffer 2.1 was added, along with 100 units of MscI and 100 units of AseI (NEB #R0534M and #R0526M) to a total volume of 370 μl in a 1.5 ml lo-bind Eppendorf tube. This was placed on a rotator at 12 rpm at 37 °C overnight. DNA concentration was then quantified using a Qubit Broad Range DNA kit (Thermo Fisher #Q32850). DNA was then mixed with orange loading buffer and loaded on a 0.3% TAE agarose gel made with Lonza SeaKem Gold agarose (# 50512) and 15 μl SYBRSafe gel stain (Thermo Fisher #S33102) per 100 ml gel. A GeneRuler High Range DNA Ladder (Thermo Fisher SM1351) was loaded in an adjacent lane. To avoid overloading, DNA was loaded with no more than 250 ng per mm of lane width (~30μg per sample). The gel was run at 2 V/cm for 1 hour and imaged over a blue light transilluminator. The gel was cut to remove fragments smaller than 20 kb, while keeping everything larger, up to the well itself. DNA was purified from the resulting gel slice using a Zymoclean Large Fragment DNA Recovery Kit (Zymo # D4045), with modifications: the gel slice was melted at room temperature on a rotator at 12 rpm, and DNA was eluted from the column twice with elution buffer heated to 70 °C. The DNA was then quantified by Qubit again. DNA was prepared for sequencing using an ONT LSK-109 native library prep kit, and sequenced on a v9.4 MinION flow cell. CENP-A targeted DiMeLo-seq was performed on unfixed HG002 cells processed in parallel with IgG targeted, free-floating pA-Hia5, and untreated samples. For each treatment ~25 million cells were processed in 5 tubes of ~5 million cells each. DiMeLo-seq was initially performed as described above. Alpha-HOR-RES was performed on these samples and 250ng-1ug of recovered DNA of each sample was then processed for Nanopore sequencing using method (1), described above.

### Basecalling, modification calling, and LMNB1 data analysis

All sequencing was performed on ONT MinION v9.4 flow cells. Basecalling and modification calling were performed on Amazon Web Services g4dn.metal instances, which have 8 NVIDIA T4 GPUs, 96 CPUs, 384 Gb memory, and 2×900 Gb local solid-state storage; this configuration allows for efficient parallelization and high basecalling speed. Basecalling was first performed using Oxford Nanopore Technologies’s Guppy software (v4.5.4), using a Rerio res_dna_r941_min_modbases-all-context_v001.cfg basecalling model, and demultiplexing when appropriate. Modification calls were extracted from fast5 output files using ont-pyguppy-client-api. Basecalled reads were aligned to the T2T-CHM13v1.0 reference sequence using Winnowmap (v2.03), which is adapted to perform better than other long-read aligners in repetitive regions (Jain et al. 2020). Fast5 files were split by barcode using fast5_subset then re-basecalled using ONT’s Megalodon software (v2.3.1), using the same reference and model file. Custom code was used to parse output files and will be made available on Github. To evaluate performance, cLAD and ciLAD coordinates (Altemose et al. 2020) were lifted over from hg38 to the T2T-CHM13v1.0 reference (Nurk et al. 2021). A read was assigned to a cLAD or ciLAD bin if it overlapped the bin with more than 50% of its length, and any mA calls on that read were assigned to that bin for browser plotting. Single-cell Dam-LMNB1 data were re-mapped to T2T-CHM13v1.0 and processed as described in (Altemose et al. 2020). For plotting Fig. 3d, 100 kb bins were filtered to those with at least 60 overlapping DiMeLo-seq reads, and with a single-cell combined mean-squared-error estimate <0.004, to select for regions with higher-confidence contact frequency estimates. Browser plots were made using the WashU Epigenome Browser (D. Li et al. 2019). Custom scripts are available by request and will be available on github upon publication.

### CTCF data analysis

For all GM12878 samples (CTCF-targeted, IgG control, free pA-Hia5, in vitro methylated genomic DNA, and untreated genomic DNA), Megalodon modified basecalls were used for analysis. Reference GM12878 ChIP-seq peaks (ENCFF797SDL, (ENCODE Project Consortium 2012)) were lifted over from hg38 to T2T chm13v1.0. These peaks were intersected with known CTCF motifs that were also lifted over to T2T chm13v1.0 (Kheradpour and Kellis 2014). Enrichment in CTCF ChIP-seq peaks was calculated using bedtools (v2.28.0). The anti-CTCF antibody (ab188408) was confirmed by personal correspondence to bind a peptide in the first 600 C-terminal amino acids of the protein.

For analysis of CTCF-targeted DiMeLo-seq data, modified basecalls for reads spanning CTCF ChIP-seq motifs were extracted within −1 kb to 1 kb of the motif center. To extract single molecules spanning peaks, pysam (v0.15.3) was used with code adapted from (De Coster, Stovner, and Strazisar 2020). If the motif was on the - strand, the positions of bases relative to the motif center were flipped. Only non-overlapping CTCF sites within the −1 kb to 1 kb display were considered, and only mA called with probability > 0.9 were plotted. Aggregate profiles were plotted with a moving average of 50 bp. For peak and read counts considered, see Supplementary Figure 4d. For joint analysis of mA and mCpG on the same molecules, only molecules spanning motifs in the upper quartile of ChIP-seq peaks that have at least one mA called with probability > 0.9 and one mCpG called with probability > 0.9 were considered, resulting in 5,054 reads considered.

To call CTCF peaks on single molecules, all molecules spanning quartile 4 ChIP-seq peaks that had at least one mA detected with probability > 0.9 were considered. Reads were filtered to require they span the CTCF motif with at least 100 bp covered on each side of the motif. With a sliding window of 20 A’s the probability that at least one A was methylated within the bin was computed by calculating 1-exp(sum(log(1-p))) for each mA with probability > 0.5. Reads that had at least one binned probability > 0.9 have a called peak, resulting in 10,631 reads with called peaks. The peak center was calculated as the midpoint of the longest stretch of mA with binned probability > (2% FDR). The FDR was calculated as the fraction of adenines methylated in the unmethylated control divided by the fraction of adenines methylated in the CTCF-targeted sample. The distribution of differences between predicted peak center and known motif center was fit using scipy (v1.4.1) to a Student’s t-distribution with 0.86 degrees of freedom (fit parameters were df: 0.86, loc: −7.87 scale: 89.7). The interval which contains 70% of the distribution was reported for the single-molecule resolution. To calculate the FDR for the single-molecule sensitivity for CTCF, we computed the fraction of molecules containing a peak in the IgG control divided by the fraction of molecules containing a peak in the CTCF-targeted sample for the top decile of ChIP-seq peaks.

For analysis of single molecules spanning two CTCF sites, peak pairs that were 2 to 10 kb apart were selected from the strongest CTCF ChIP-seq peaks (quartiles 3 and 4). As in peak calling, binned qualities in bins of 20 A’s were computed. Here, if a binned probability > 0.9 fell within 100 bp on either side of at least one of the two CTCF binding sites, the read was considered to have a correctly called peak and the molecule was included in Figure 4d. A total of 625 peak pairs were considered with a total of 511 reads spanning these peaks. Reads were clustered using kmeans clustering (scikit-learn v0.24.2) with 4 clusters, and the reads within clusters were sorted by binned mA signal within 100 bp of the left peak.

### H3K9me3 data analysis

For all HG002 samples (H3K9me3-targeted, IgG control, and free pA-Hia5) a merged bam file was created with samtools (v1.8) from the Guppy bam and winnowmap outputs aligned to a special male reference genome (CHM13+HG002X+hg38Y: autosomes from the T2T chm13v1.0 genome combined with a T2T assembly of HG002 chromosome X (Nurk et al. 2021) and the chrY sequence from hg38), and a mapping quality threshold of 10 was applied. To compare to CUT&RUN, broad peaks were called using macs2 (v2.1.1) on a H3K9me3 CUT&RUN bam file from HG002 (Altemose et al. 2021). Centromere and HOR boundaries were defined from the T2T centromere annotation (Altemose et al. 2021). Enrichment in CUT&RUN peaks, centromeres, and active HOR arrays was computed using bedtools (v2.28.0). For analyzing mA signal at HOR boundaries, the mean mA/A in a 60 kb rolling window from −300 kb within the HOR to 300 kb outside of the HOR was computed. A total of 2,359 reads spanned this region. HOR boundaries considered were those that transition quickly into non-repetitive sequences: 1p, 2pq, 6p, 9p, 13q, 14q, 15q, 16p, 17pq, 18pq, 20p, 21q, 22q. For single molecule browser visualization, modified bases were extracted as in CTCF analysis using custom python scripts, and modified bases with probability > 0.9 were displayed. Single-molecule browser plots were generated using plotly (v4.5.2).

### CENP-A data analysis

Basecalling for centromere enriched samples was performed twice both times using Guppy (5.0.7). The first basecalling used the “super accuracy” basecalling model (dna_r9.4.1_450bps_sup.cfg), followed by alignment to the CHM13+HG002X+hg38Y reference genome using Winnowmap (v2.03). These alignments were then filtered for only primary alignments and mapq score greater than 10 using samtools view -F 2308 -q10. A second round of basecalling was then performed again using Guppy (5.0.7) but now with the rerio all-context basecalling model (res_dna_r941_min_modbases-all-context_v001.cfg) with --bam_out and -- bam_methylation_threshold 0.0. Modified basecalls were then merged by read id with winnowmap alignments to generate bam files with high confidence alignments combined with modification calls for downstream processing. Two independent biological replicates, the first which was sequenced on two separate flow cells, for three total sequencing runs were merged for the final analysis.

To calculate centromere enrichment samtools bedcov was used to calculate the total bases that mapped to alpha satellite active HORs in free-floating Hia5 treated samples treated with and without centromere enrichment (Danecek et al. 2021). Reported coverage at each centromeric region is relative to the length of that region. Chromosomes with more than one active HOR had the mean value of length-normalized coverage reported. deepTools2 bamCoverage (v3.3.1) (Ramírez et al. 2016) was used to generate bigWigs with 10 kb bin size, that were plotted on HG002 chromosome X using pygenometracks (v3.6) (Lopez-Delisle et al. 2021) to compare chromosome-wide coverage between centromere enriched and unenriched samples.

A *k*-mer counting pipeline was used to identify CENP-A enriched *k*-mers from chm13 Native ChIP-seq experiment (Logsdon et al. 2021; Smith et al. 2021). After separating DiMeLo-seq reads into those that did and did not have a CENP-A enriched *k*-mer, methylation frequency for each subset was calculated, as well as the fold enrichment for percentage mA/A of reads containing CENP-A enriched *k*-mers over those that did not.

For single molecule browser visualization, modified bases were extracted as in CTCF analysis using custom python scripts, and modified bases with mA probability (from Guppy) > 0.9 and mCpG probability (from Guppy) > 0.8 were displayed.

Average fraction of mA or mCpG methylation for aggregate views were calculated as the fraction of reads at each adenine that have probability score (from Guppy) greater than 0.9 for mA or 0.8 for mCpG. Representative plots show average fraction of reads at each adenine or CpG with methylation probability score above threshold binned by smoothing over a rolling window of 200 bp (for plots showing mA or mCpG within 50 - 60 kb regions) or 1 kb (for plot showing mA over 200 kb region of Chromosome X). Coverage plots indicate the number of reads that are aligned with each adenine position within the region.

For estimating nucleosome density per read, the number of nucleosomes on each read was estimated within incremental 5 kb sliding windows (step size of 2500 bp) as follows. Within each 5 kb window, all sliding 500 bp bins (step size of 1 bp) with > 0.9 probability of containing at least 1 mA were identified on each read and contiguous bins were counted as one nucleosome. The highest probability of directed methylation was assumed to be within ~160 bp on either side of a nucleosome that wraps ~175 bp. Average number of nucleosomes per read was calculated by only considering reads that align to at least 1 kb within each 5 kb window.

