## Supplementary Material for "DiMeLo-seq: a long-read, single-molecule method for mapping protein-DNA interactions genome-wide"

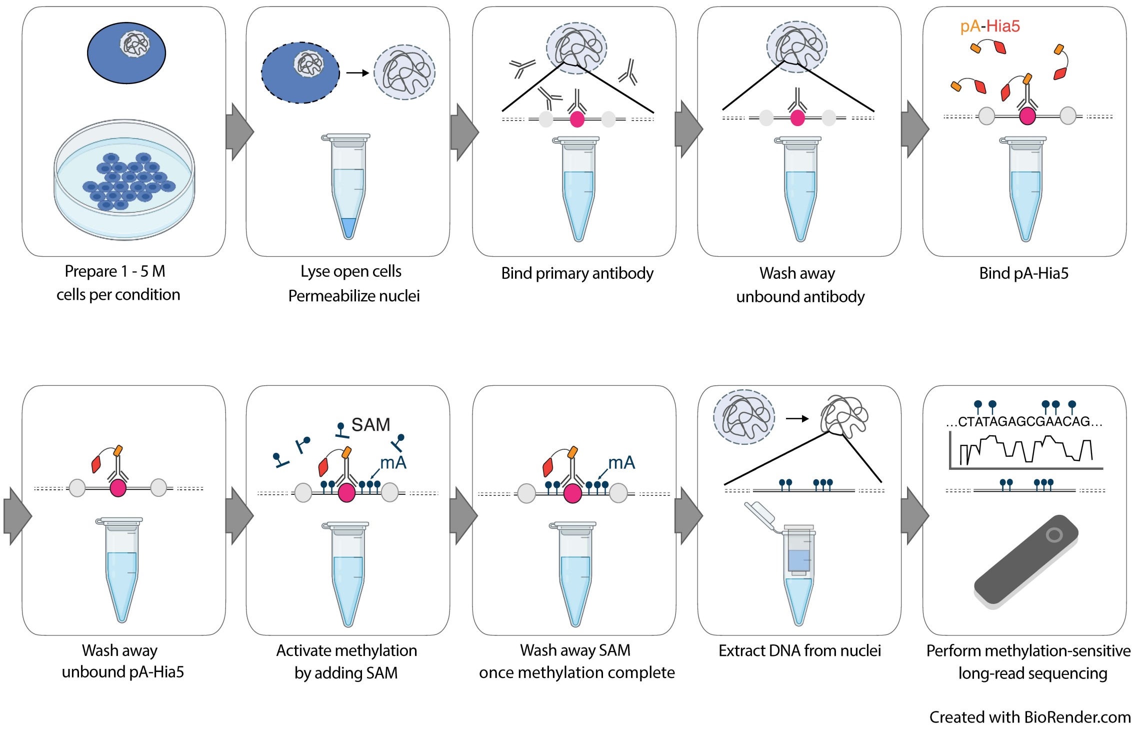


**Supplementary Figure 1. Workflow of DiMeLo-seq *in situ* methylation, DNA extraction, and sequencing**. Schematic of the DiMeLo-seq *in situ* methylation protocol, which involves a series of binding steps and washes followed by DNA extraction and sequencing.

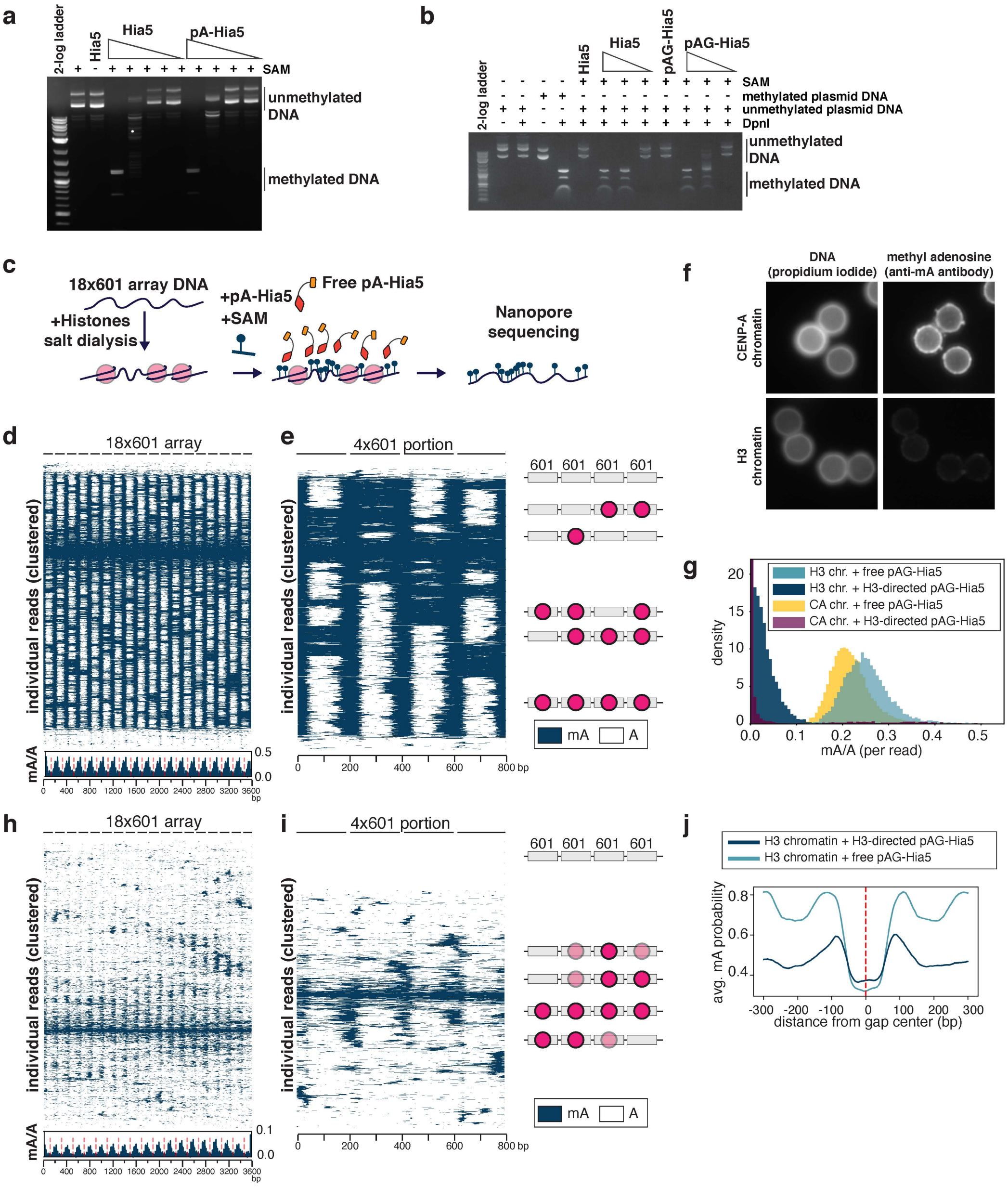


**Supplementary Figure 2. In vitro assessment of methylation of DNA and chromatin by pA-Hia5 and pAG-Hia5. a,b,** Agarose gel electrophoresis image of DpnI digestion of (unmethylated) plasmid DNA following incubation with Hia5, pA-Hia5 (a), or pAG-Hia5 (b). **c,** Schematic of methylation of artificial chromatin co-incubated with free pA-Hia5 and SAM depicting methylation of all accessible DNA. **d,e,** Heatmap showing methylation on 2000 individual reads from CENP-A chromatin following incubation with free pA-Hia5, hierarchically clustered by jaccard distance over the entire 18x601 array (d) or a subset 4x601 region along with cartoons depicting predicted nucleosome positions (e). Inset in d. shows average mA/A on every base position of 18x601 array. (red dashed line indicates 601 dyad position). **f,** Representative immunofluorescence images of chromatin-coated beads following methylation using CENP-A-directed pA-Hia5. **g,** Histogram of fraction of methylation (mA/A) on reads from CENP-A or H3 chromatin methylated with free pAG-Hia5 or H3-directed pAG-Hia5. **h,i.** Heatmap showing methylation on 2000 individual reads from H3 chromatin following H3-directed methylation, hierarchically clustered by jaccard distance over the entire 18x601 array (h) or a subset 4x601 region (i). Inset in (h). shows average mA/A on every base position of the 18x601 array. (red dashed line indicates 601 dyad position). **j,** Average methylation probability score (from Megalodon base-calling) near the center of 100-180 bp gaps (red dashed line) in mA signal on reads from CENP-A chromatin methylated using free pAG-Hia5 incubation or H3-directed pA-Hia5.

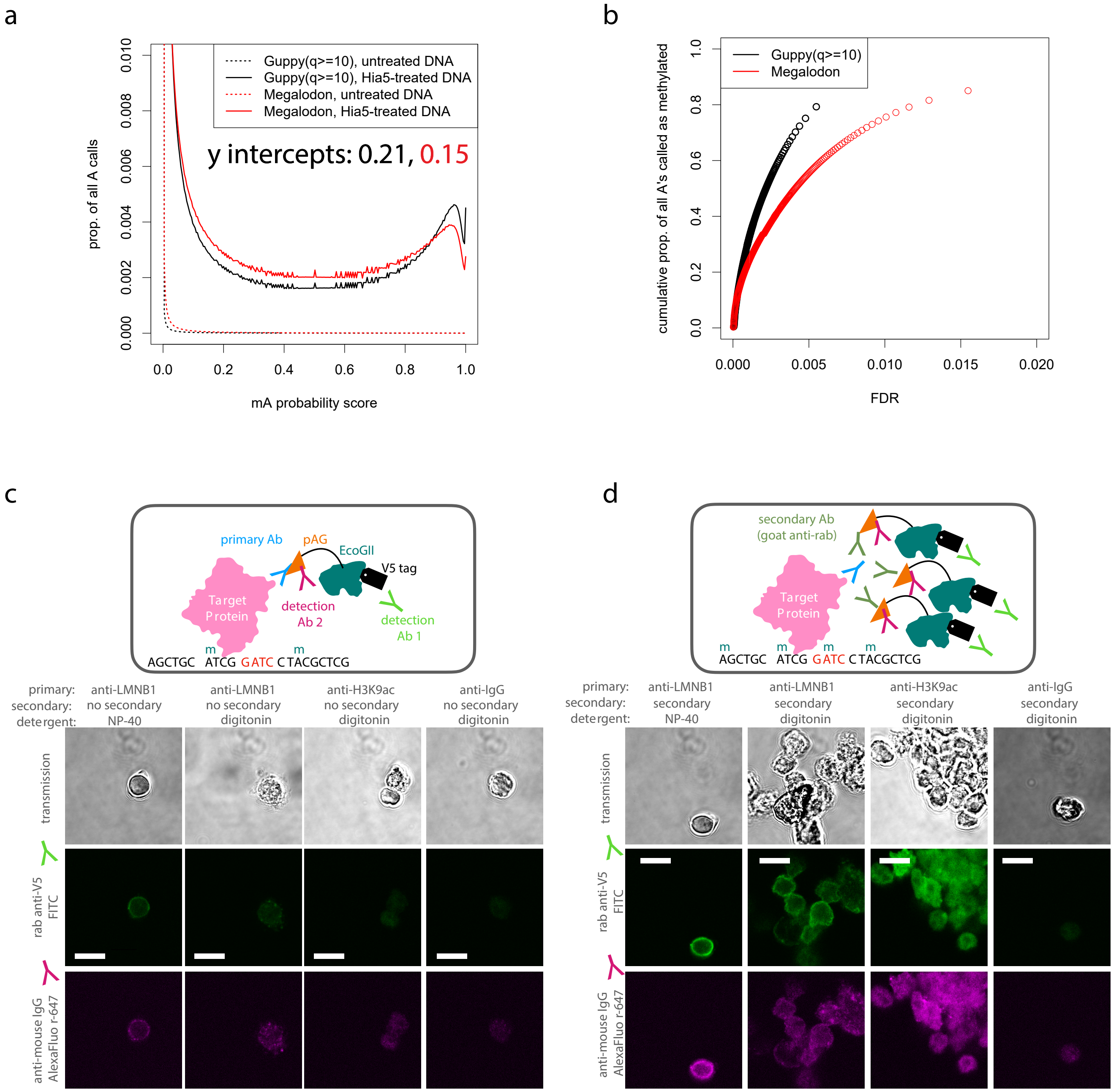


**Supplementary Figure 3. Assessment of mA calling and LMNB1 targeting. a,** The proportion of all adenines called as methylated at each possible probability threshold using two different software packages on ONT reads from two HEK293T DNA samples: untreated genomic DNA and naked genomic DNA methylated by Hia5 *in vitro*. The untreated DNA provides a measure of the false positive rate at each threshold, since it contains few or no methyl adenines. **b,** Estimates of the proportion of As methylated in the Hia5-treated DNA sample at each FDR threshold (determined from a). Roughly 80% of the adenines on the Hia5-treated DNA appear to be methylated. **c-d,** In the DiMeLo-seq workflow, following the primary antibody and pA/G-MTase binding and wash steps, a sample of nuclei can be taken for quality assessment by immunofluorescence. One can determine the locations and relative quantity of pA/G-MTase molecules using fluorophore-conjugated antibodies that bind to the pA/G-MTase but not to the primary antibody. In these images, the results for pAG-EcoGII are shown, comparing different antibodies, detergents, and samples with (d) and without (c) the use of an unconjugated secondary antibody to recruit more pA/G-MTase molecules to the target protein. Scale bar: 10 microns.

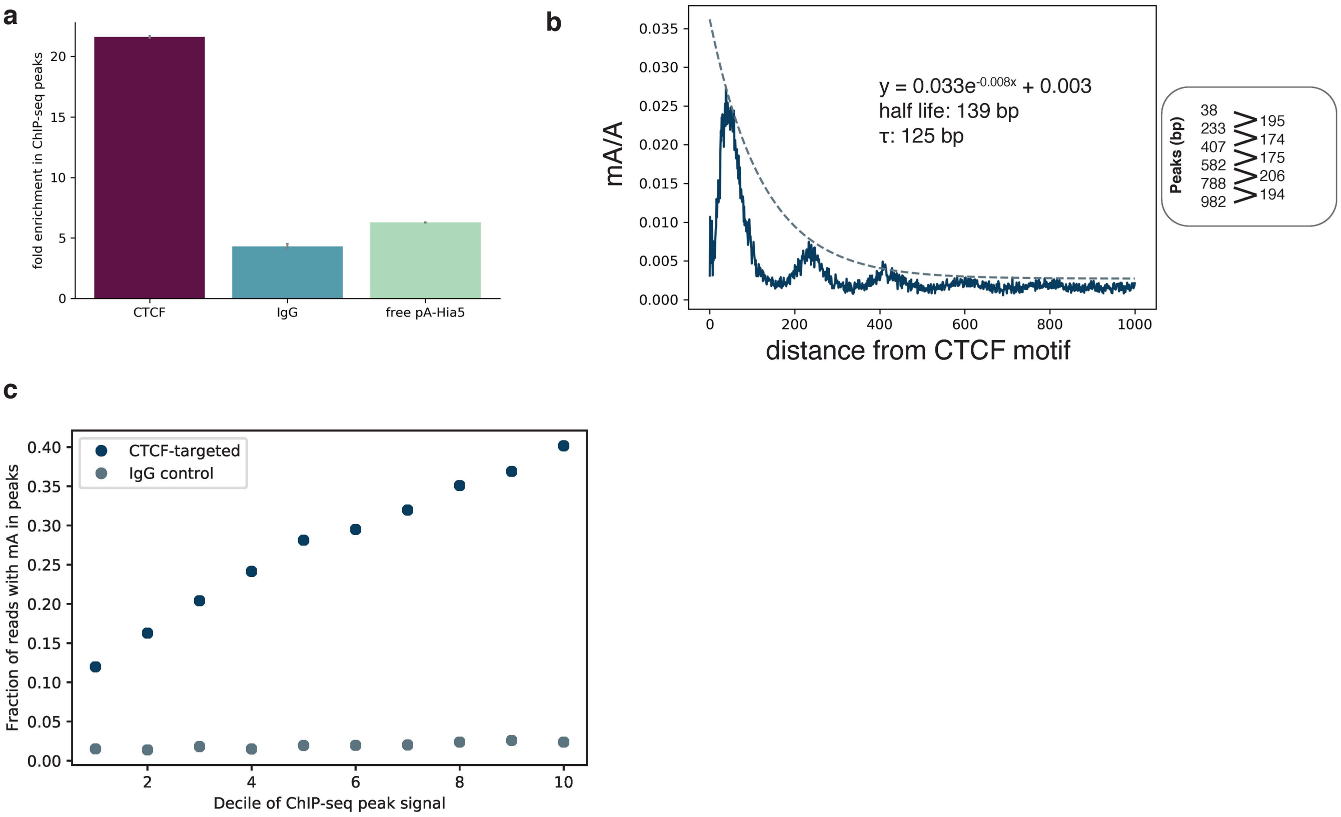


d

| Quartile | Number of peaks | Number of reads | Number of reads with at least one mA >thresh90 | Number of reads with at least one mCpG >thresh90 |
|---|---|---|---|---|
| 4 | 4812 | 27204 | 10486 | 10524 |
| 3 | 4811 | 28155 | 9079 | 9812 |
| 2 | 4811 | 27673 | 7500 | 8990 |
| 1 | 4812 | 27350 | 5154 | 9633 |

**Supplementary Figure 4. Analysis of specificity, methylation decay rate, and sensitivity in CTCF targeting a,** Fold enrichment over background of mA/A in ChIP-seq peak regions. Error bars represent the 95% confidence interval determined by simulating the proportion of adenines methylated as a Beta distribution with a uniform prior. **b,** Methylation decay from the CTCF motif center for the upper quartile of ChIP-seq signal is fit with an exponential decay function. The positions of the peaks are indicated, with the spacing between peaks also noted. **c,** Fraction of reads that contain at least one mA called with probability > 0.9 within 100 bp on either side of the motif center for each decile of ChIP-seq peak strength for the CTCF-targeted sample and IgG control. **d,** Number of peaks and reads displayed in Figure 4a.

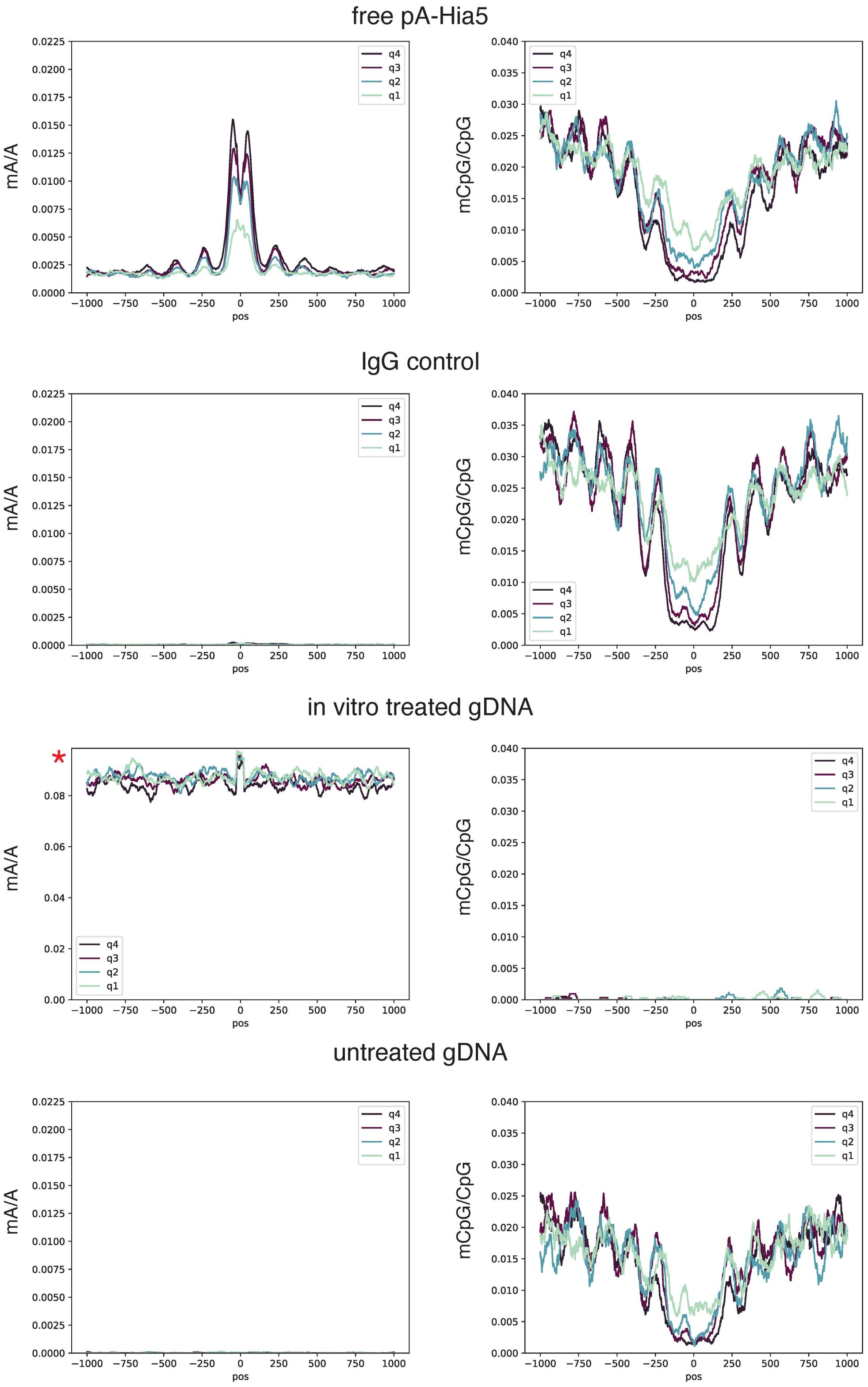


**Supplementary Figure 5. Control mA and mCpG profiles at CTCF peaks.** Profiles at CTCF ChIP-seq peaks for free pA-Hia5, IgG control, *in vitro* treated genomic DNA, and untreated genomic DNA. Quartiles indicate rank of ChIP-seq peak strength. All axes are the same scaling as in Figure 4, except for mA/A of *in vitro* treated gDNA. With high mA levels achieved only with this *in vitro* methylated control, mC basecalling fails.

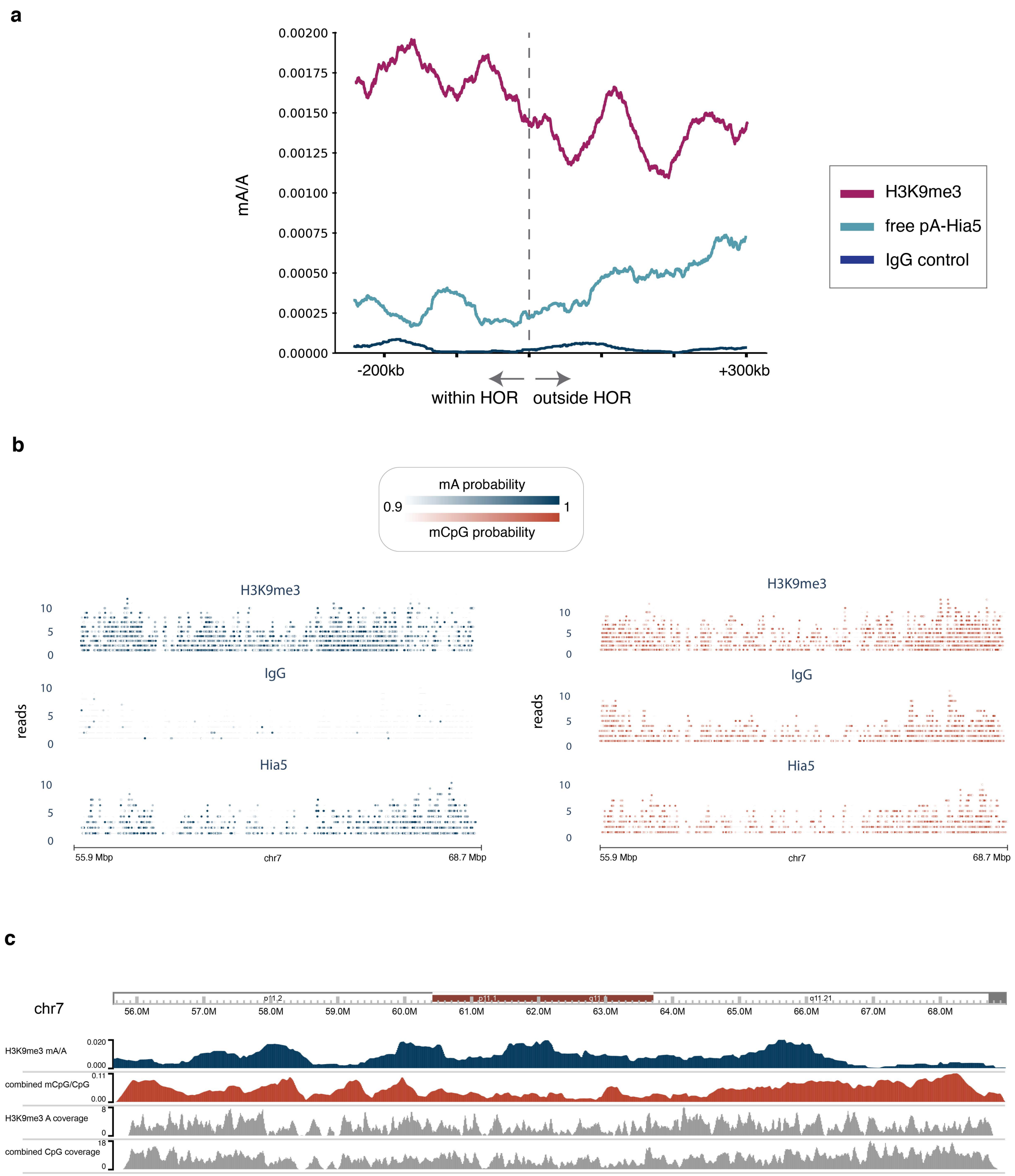


**Supplementary Figure 6. H3K9me3 control analysis at HOR boundaries and in centromere 7. a,** Density of methylated adenines for the H3K9me3-targeted sample and IgG and free pA-Hia5 controls in 60 kb sliding window across HOR boundaries 1p, 2pq, 6p, 9p, 13q, 14q, 15q, 16p, 17pq, 18pq, 20p, 21q, 22q. **b,** Centromere 7 single molecule browser tracks for H3K9me3-targeted sample, IgG control, and free pA-Hia5. The same molecules are shown in both plots, with mA calls indicated in the first, and mCpG calls indicated in the second. **c,** Coverage tracks in 10 kb bins to accompany mA/A and mCpG/CpG tracks from Figure 5d.

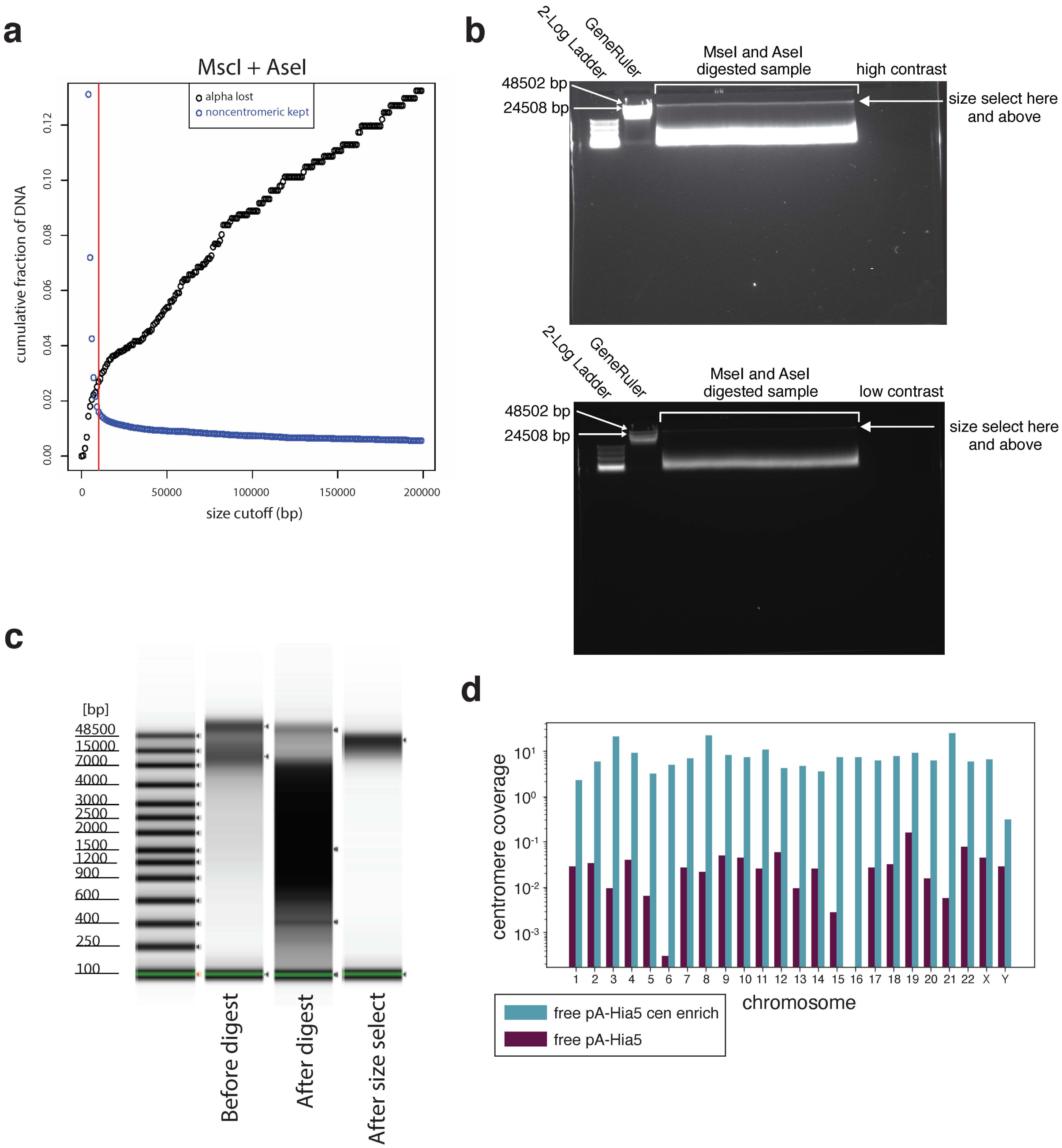


**Supplementary Figure 7. AlphaHOR-RES centromere enrichment design and implementation**

**a,** Simulated cumulative distribution of the proportion of alpha-satellite DNA lost (black) and non-centromeric DNA kept (blue) after MscI and AseI digestion of the T2T chm13 genome at different size selection cutoffs. **b,** Representative image of agarose gel run on total genomic DNA after Msc1 and Ase1 digestion. Left lane is 2-log ladder and the next lane is GeneRuler High Range DNA ladder, followed by digested sample. Arrow indicates the location where gel is cut. Sample recovered from above cut site. Top is high contrast, bottom is reduced contrast. **c,** genomic DNA tapestation gel images of sample before digestion (left lane), after digestion (middle lane), and after size selection (right lane). **d,** Coverage of the active HOR on each chromosome from the CHM13+HG002X+hg38Y reference genome from free floating pA-Hia5 DiMeLo-seq libraries that did and did not have alphaHOR-RES performed.

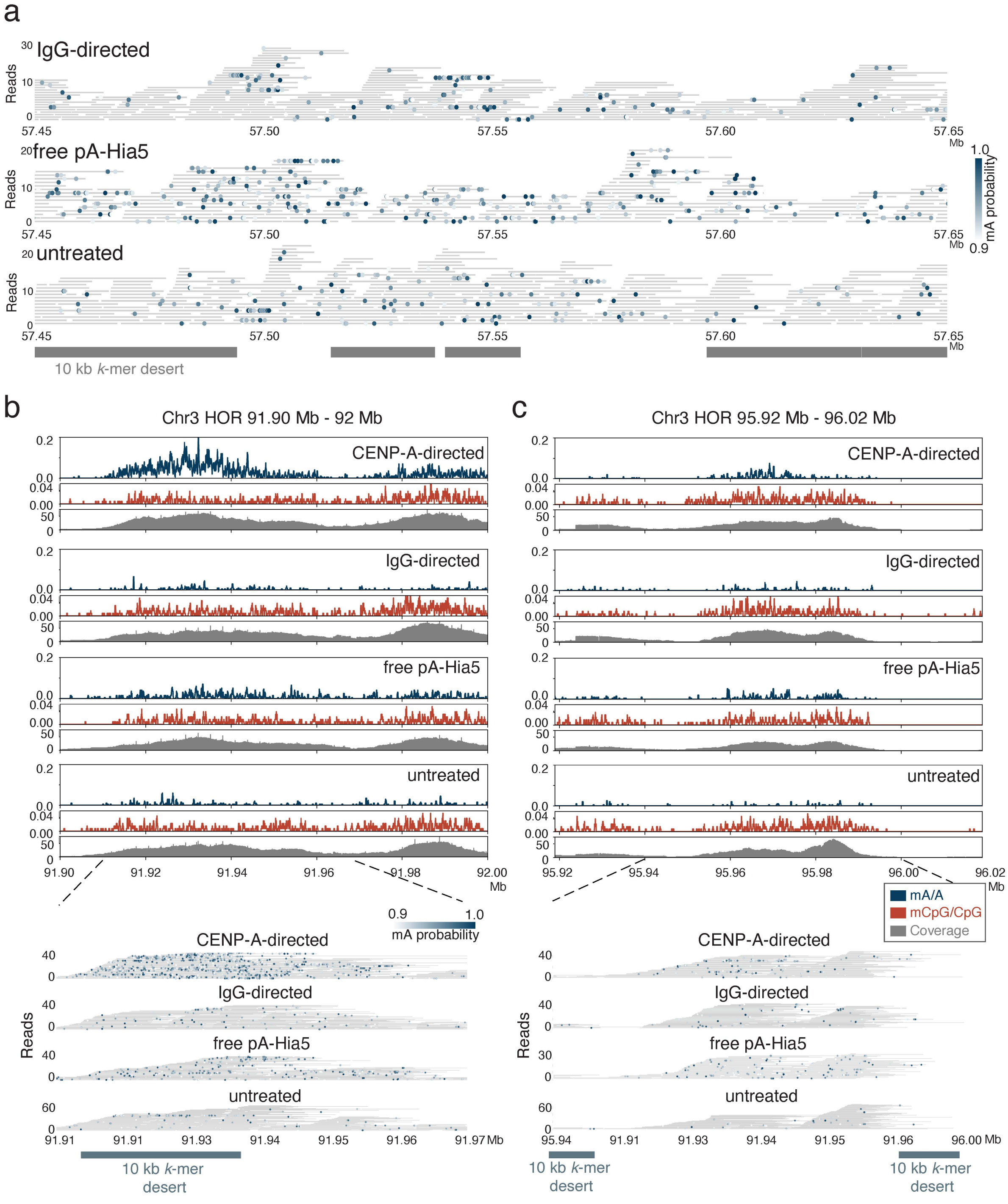


**Supplementary Figure 8. Methylation within higher order repeats on chromosomes X and 3. a,** Single molecule view with individual reads in gray and mA depicted as dots for IgG targeted DiMeLo-seq libraries or free pA-Hia5 treated or untreated libraries. Scale bar indicates the probability of adenine methylation (from Guppy) between 0.9 and 1. Regions with at least 10 kb without unique 51 bp *k*-mers shown in grey to illustrate difficult to map locations for short-read sequencing. **b,** Aggregate mA and mCpG signal on chromosome 3 HOR between 91.9 and 92.0 Mb for all conditions. mA/A and mCpG/CpG plots indicate fraction of reads with methylation above threshold (0.9 for mA, 0.8 for mCpG) probability (average over 200 bp rolling window for visualization). Below are single molecule views with individual reads in gray and mA depicted as dots for CENP-A targeted, IgG targeted DiMeLo-seq libraries or free pA-Hia5 treated or untreated libraries. Scale bar indicates the probability of adenine methylation (from Guppy) between 0.9 and 1. Regions with at least 10 kb without unique 51 bp *k*-mers shown in grey to illustrate difficult to map locations for short-read sequencing. **c,** Same as b, for distinct chromosome 3 HOR located between 95.92 and 96.02 Mb. Scale bars indicate the probability of adenine methylation (from Guppy) between 0.9 and 1.

| ID | Batch | BC | Cell Line | Ab | Ab dil. | Other / Notes | pA/G | Link. len. (aa) | MTase | [MTase] (nM) | Ab2 | pA/G bind temp | Act. buf. | Act. [SAM] (uM) | Read number | Total bases sequenced | Mean read len. | ON:OFF | ON-target prop. mA | All reads prop. mA |
|---|---|---|---|---|---|---|---|---|---|---|---|---|---|---|---|---|---|---|---|---|
| **1** | *1* | *6* | HEK293T | LMNB1 | 500 | SRE XL | pAG | 29 | EcoGII | 50 | N | 4C | A | 500 | 86,481 | 1,976,058,348 | 22,850 | 1.668 | 4.46E-05 | 3.37E-05 |
| **2** | *2* | *9* | HEK293T | LMNB1 | 500 | SRE XL, Ab2 bind at 4C | pAG | 29 | EcoGII | 150 | Y | 4C | A | 500 | 38,539 | 847,065,308 | 21,979 | 2.929 | 1.57E-04 | 1.00E-04 |
| **3** | *2* | *10* | HEK293T | LMNB1 | 500 | SRE XL | pAG | 29 | EcoGII | 150 | N | 4C | A | 500 | 36,921 | 856,905,359 | 23,209 | 2.499 | 1.37E-04 | 8.96E-05 |
| **4** | *2* | *14* | HEK293T | LMNB1 | 500 | NP40 0.5%,SRE XL,Ab2@4C | pAG | 29 | EcoGII | 150 | Y | 4C | A | 500 | 60,602 | 1,490,024,882 | 24,587 | 1.208 | 1.18E-04 | 1.04E-04 |
| **5** | *2* | *15* | HEK293T | LMNB1 | 500 | NP40 0.5%, SRE XL | pAG | 29 | EcoGII | 150 | N | 4C | A | 500 | 38,772 | 943,586,111 | 24,337 | 1.351 | 7.23E-05 | 5.95E-05 |
| **6** | *3* | *16* | HEK293T | LMNB1 | 500 | | pAG | 29 | EcoGII | 527 | N | RT | A | 500 | 162,252 | 1,752,177,765 | 10,799 | 1.377 | 5.32E-05 | 4.83E-05 |
| **7** | *3* | *17* | HEK293T | LMNB1 | 500 | | pAG | 29 | EcoGII | 527 | Y | RT | A | 500 | 198,264 | 2,102,108,611 | 10,603 | 3.034 | 9.52E-05 | 6.15E-05 |
| **8** | *3* | *18* | HEK293T | LMNB1 | 500 | | pAG | 29 | EcoGII | 150 | N | RT | A | 500 | 106,892 | 1,227,114,851 | 11,480 | 2.046 | 2.99E-05 | 2.36E-05 |
| **9** | *3* | *19* | HEK293T | LMNB1 | 100 | | pAG | 29 | EcoGII | 527 | N | RT | A | 500 | 238,437 | 2,415,676,861 | 10,131 | 4.795 | 1.74E-04 | 9.03E-05 |
| **10** | *4* | *20* | HEK293T | LMNB1 | 100 | | pAG | 29 | EcoGII | 527 | Y | 4C | A | 500 | 121,419 | 1,261,478,986 | 10,389 | 4.729 | 4.01E-04 | 2.06E-04 |
| **11** | *4* | *21* | HEK293T | LMNB1 | 100 | | pAG | 29 | EcoGII | 527 | N | 4C | A | 500 | 123,908 | 1,276,684,561 | 10,303 | 3.982 | 2.99E-04 | 1.65E-04 |
| **12** | *4* | *23* | HEK293T | LMNB1 | 100 | 60m act. | pAG | 29 | EcoGII | 527 | N | 4C | A | 500 | 112,198 | 1,162,823,359 | 10,364 | 3.728 | 3.23E-04 | 1.87E-04 |
| **13** | *4* | *24* | HEK293T | LMNB1 | 100 | 60m act., SAM replenished | pAG | 29 | EcoGII | 527 | N | 4C | A | 500 | 119,153 | 1,207,239,004 | 10,132 | 3.049 | 3.13E-04 | 1.89E-04 |
| **14** | *4* | *1* | HEK293T | LMNB1 | 100 | 60m act., SAM replenished | pAG | 29 | EcoGII | 527 | N | 4C | A | 500 | 96,385 | 1,076,704,141 | 11,171 | 2.650 | 2.87E-04 | 1.73E-04 |
| **15** | *5* | *4* | HEK293T | LMNB1 | 100 | | pAG | 29 | EcoGII | 527 | Y | RT | A | 500 | 80,135 | 839,079,448 | 10,471 | 3.564 | 4.22E-04 | 2.36E-04 |
| **16** | *5* | *5* | HEK293T | LMNB1 | 100 | 15m act | pAG | 29 | EcoGII | 527 | Y | RT | A | 500 | 101,911 | 1,067,611,582 | 10,476 | 3.872 | 3.25E-04 | 1.74E-04 |
| **17** | *5* | *6* | HEK293T | LMNB1 | 100 | 30C act | pAG | 29 | EcoGII | 527 | Y | RT | A | 500 | 68,779 | 651,361,202 | 9,470 | 7.536 | 4.95E-04 | 2.13E-04 |
| **18** | *5* | *7* | HEK293T | LMNB1 | 100 | | pAG | 29 | EcoGII | 527 | Y | RT | A | 500 | 83,487 | 902,381,502 | 10,809 | 4.854 | 4.40E-04 | 2.35E-04 |
| **19** | *5* | *8* | HEK293T | LMNB1 | 100 | | pA | 7 | Hia5 | 527 | Y | RT | A | 500 | 26,351 | 227,185,814 | 8,622 | 5.641 | 7.04E-05 | 2.87E-05 |
| **20** | *5* | *9* | HEK293T | LMNB1 | 100 | | pA | 26 | Hia5 | 527 | Y | RT | A | 500 | 19,112 | 169,551,133 | 8,871 | 7.415 | 8.39E-05 | 4.49E-05 |
| **21** | *5* | *10* | HEK293T | LMNB1 | 100 | | pA | mix | Hia5 | 527 | Y | RT | A | 500 | 15,778 | 151,798,963 | 9,621 | 9.844 | 7.88E-05 | 3.82E-05 |
| **22** | *5* | *11* | HEK293T | LMNB1 | 100 | pAG-EcoGII+pA-Hia5-both | mix | mix | EcoGII+ | 527 | Y | RT | A | 500 | 15,975 | 149,309,499 | 9,346 | 11.823 | 2.65E-04 | 1.45E-04 |
| **23** | *5* | *12* | HEK293T | LMNB1 | 100 | NP40 0.1% | pAG | 29 | EcoGII | 527 | Y | RT | A | 500 | 86,033 | 860,416,335 | 10,001 | 2.479 | 2.50E-04 | 1.65E-04 |
| **24** | *5* | *14* | HEK293T | LMNB1 | 100 | | pAG | 29 | EcoGII | 527 | N | RT | A | 500 | 101,662 | 993,731,709 | 9,775 | 2.513 | 2.97E-04 | 1.98E-04 |
| **25** | *5* | *15* | HEK293T | LMNB1 | 100 | | pA | 7 | Hia5 | 527 | N | RT | A | 500 | 106,830 | 985,205,512 | 9,222 | 12.427 | 7.89E-04 | 3.26E-04 |
| **26** | *5* | *16* | HEK293T | LMNB1 | 100 | | pA | 26 | Hia5 | 527 | N | RT | A | 500 | 74,873 | 793,137,043 | 10,593 | 11.724 | 1.06E-03 | 4.23E-04 |
| **27** | *5* | *17* | HEK293T | LMNB1 | 100 | | pA | mix | Hia5 | 527 | N | RT | A | 500 | 93,269 | 988,096,097 | 10,594 | 17.681 | 8.36E-04 | 3.43E-04 |
| **28** | *5* | *18* | HEK293T | LMNB1 | 100 | pAG-EcoGII+pA-Hia5-both | mix | mix | EcoGII+ | 527 | N | RT | A | 500 | 92,725 | 945,637,416 | 10,198 | 6.085 | 7.71E-04 | 3.27E-04 |
| **29** | *5* | *19* | HEK293T | LMNB1 | 100 | **light fixation** | pAG | 29 | EcoGII | 527 | Y | RT | A | 500 | 91,520 | 686,940,638 | 7,506 | 5.370 | 4.31E-04 | 2.16E-04 |
| **30** | *6* | *21* | HEK293T | LMNB1 | 100 | Triton 0.1% | pA | 26 | Hia5 | 527 | N | RT | A | 500 | 100,177 | 1,126,513,813 | 11,245 | 4.664 | 9.10E-05 | 4.20E-05 |
| **31** | *6* | *22* | HEK293T | LMNB1 | 100 | Triton 0.1% | pA | 26 | Hia5 | 527 | N | RT | B | 500 | 114,462 | 1,105,844,974 | 9,661 | 21.179 | 3.68E-04 | 1.36E-04 |
| **32** | *6* | *23* | HEK293T | LMNB1 | 100 | NP40 0.1% | pA | 26 | Hia5 | 527 | N | RT | A | 500 | 105,547 | 934,810,760 | 8,857 | 3.679 | 6.12E-05 | 2.82E-05 |
| **33** | *6* | *24* | HEK293T | LMNB1 | 100 | NP40 0.1% | pAG | 29 | EcoGII | 527 | N | RT | A | 500 | 84,310 | 868,314,441 | 10,299 | 4.312 | 1.11E-04 | 5.22E-05 |
| **34** | *6* | *1* | HEK293T | LMNB1 | 100 | | pAG | 29 | EcoGII | 527 | N | RT | A | 500 | 91,978 | 1,189,823,234 | 12,936 | 3.411 | 1.49E-04 | 7.60E-05 |
| **35** | *6* | *2* | HEK293T | LMNB1 | 100 | | pA | 26 | Hia5 | 527 | N | RT | A | 500 | 105,110 | 1,264,716,882 | 12,032 | 24.212 | 5.33E-04 | 1.78E-04 |
| **36** | *6* | *3* | HEK293T | LMNB1 | 100 | | pA | 26 | Hia5 | 527 | N | RT | B | 500 | 103,978 | 1,311,352,305 | 12,612 | 38.048 | 2.06E-03 | 6.93E-04 |
| **37** | *7* | *4* | HEK293T | LMNB1 | 100 | | pA | 26 | Hia5 | 527 | N | RT | B* | 500 | 90,785 | 752,583,312 | 8,290 | 26.474 | 4.08E-03 | 1.47E-03 |
| **38** | *7* | *5* | HEK293T | LMNB1 | 100 | | pAG | 7 | Hia5 | 527 | N | RT | B* | 500 | 48,122 | 459,680,632 | 9,552 | 29.831 | 2.23E-03 | 8.67E-04 |
| **39** | *7* | *6* | HEK293T | LMNB1 | 100 | | pAG | 29 | EcoGII | 527 | N | RT | B* | 500 | 81,695 | 727,342,635 | 8,903 | 7.484 | 1.21E-03 | 5.25E-04 |
| **40** | *7* | *7* | HEK293T | LMNB1 | 100 | | pA | 26 | Hia5 | 527 | N | RT | B | 500 | 65,117 | 583,081,677 | 8,954 | 23.486 | 3.05E-03 | 1.20E-03 |
| **41** | *7* | *8* | HEK293T | LMNB1 | 100 | | pAG | 7 | Hia5 | 527 | N | RT | B | 500 | 56,317 | 540,388,209 | 9,595 | 29.747 | 2.08E-03 | 8.06E-04 |
| **42** | *7* | *9* | HEK293T | LMNB1 | 100 | | pAG | 29 | EcoGII | 527 | N | RT | B | 500 | 47,684 | 449,643,915 | 9,430 | 8.223 | 1.15E-03 | 4.77E-04 |
| **43** | *7* | *10* | HEK293T | LMNB1 | 100 | 30C act | pA | 26 | Hia5 | 527 | N | RT | B | 500 | 58,489 | 523,464,949 | 8,950 | 29.977 | 2.24E-03 | 8.07E-04 |
| **44** | *7* | *11* | HEK293T | LMNB1 | 100 | 30C act | pAG | 7 | Hia5 | 527 | N | RT | B | 500 | 57,609 | 511,675,484 | 8,882 | 17.415 | 1.26E-03 | 4.72E-04 |
| **45** | *7* | *12* | HEK293T | LMNB1 | 100 | 30C act | pAG | 29 | EcoGII | 527 | N | RT | B | 500 | 61,562 | 551,698,966 | 8,962 | 4.400 | 5.33E-04 | 2.44E-04 |
| **46** | *7* | *13* | HEK293T | LMNB1 | 100 | 30C act | pAG | 29 | EcoGII | 527 | N | RT | A | 500 | 71,900 | 628,811,974 | 8,746 | 8.180 | 3.02E-04 | 1.31E-04 |
| **47** | *7* | *14* | HEK293T | LMNB1 | 100 | Ab2 (GP) | pA | 26 | Hia5 | 527 | Y | RT | B | 500 | 99,602 | 843,096,447 | 8,465 | 21.368 | 2.33E-03 | 8.98E-04 |
| **48** | *7* | *15* | HEK293T | LMNB1 | 100 | Ab2 (GP) | pA | 26 | Hia5 | 527 | Y | RT | A* | 500 | 54,677 | 484,736,060 | 8,865 | 8.614 | 4.85E-04 | 2.04E-04 |
| **49** | *7* | *16* | HEK293T | LMNB1 | 100 | Ab2 (GP) | pAG | 7 | Hia5 | 527 | Y | RT | B | 500 | 63,400 | 571,575,224 | 9,015 | 13.879 | 1.43E-03 | 5.34E-04 |
| **50** | *7* | *17* | HEK293T | LMNB1 | 100 | Ab2 (GP), 30C act | pAG | 29 | EcoGII | 527 | Y | RT | A | 500 | 60,290 | 517,001,320 | 8,575 | 7.268 | 3.43E-04 | 1.68E-04 |
| **51** | *7* | *18* | HEK293T | LMNB1 | 100 | pA-Hia5 short+long | pA | mix | Hia5 | 527 | N | RT | B | 500 | 47,923 | 468,996,420 | 9,786 | 21.990 | 3.19E-03 | 1.31E-03 |
| **52** | *7* | *19* | HEK293T | LMNB1 | 100 | pA-Hia5 200 nM, Ab2 (GP) | pA | 26 | Hia5 | 200 | Y | RT | B | 500 | 52,514 | 513,948,158 | 9,787 | 19.589 | 2.82E-03 | 1.06E-03 |
| **53** | *7* | *20* | HEK293T | LMNB1 | 100 | pA-Hia5 50 nM, Ab2 (GP) | pA | 26 | Hia5 | 50 | Y | RT | B | 500 | 48,723 | 441,526,404 | 9,062 | 20.303 | 2.55E-03 | 9.19E-04 |
| **54** | *7* | *22* | HEK293T | LMNB1 | 100 | **frozen** | pA | 26 | Hia5 | 527 | N | RT | B | 500 | 71,276 | 488,718,798 | 6,857 | 28.059 | 2.98E-03 | 1.09E-03 |
| **55** | *7* | *23* | HEK293T | LMNB1 | 100 | Ab2 (Goat), 30C act | pAG | 29 | EcoGII | 527 | Y | RT | A | 500 | 61,444 | 554,745,469 | 9,028 | 8.711 | 5.32E-04 | 2.22E-04 |
| **56** | *8* | *1* | HEK293T | LMNB1 | 100 | poor batch | pA | 7 | Hia5 | 200 | N | RT | B* | 500 | 45,362 | 345,941,926 | 7,626 | 7.676 | 2.36E-03 | 1.09E-03 |
| **57** | *8* | *2* | HEK293T | LMNB1 | 100 | poor batch | pA | 7 | Hia5 | 200 | N | RT | B* | 500 | 40,187 | 353,173,477 | 8,788 | 6.865 | 2.38E-03 | 1.06E-03 |
| **58** | *8* | *3* | HEK293T | LMNB1 | 50 | poor batch | pA | 7 | Hia5 | 200 | N | RT | B* | 500 | 33,387 | 278,128,346 | 8,330 | 9.621 | 3.97E-03 | 1.73E-03 |
| **59** | *8* | *7* | GM12878 | LMNB1 | 100 | poor batch | pA | 7 | Hia5 | 200 | N | RT | B* | 500 | 55,380 | 462,838,466 | 8,358 | 8.386 | 2.75E-03 | 1.24E-03 |
| **60** | *8* | *8* | HG002 | LMNB1 | 100 | poor batch | pA | 7 | Hia5 | 200 | N | RT | B* | 500 | 59,508 | 439,509,594 | 7,386 | 9.339 | 3.13E-03 | 1.30E-03 |
| **61** | *8* | *9* | Hap1 | LMNB1 | 100 | poor batch | pA | 7 | Hia5 | 200 | N | RT | B* | 500 | 59,648 | 424,749,575 | 7,121 | 7.473 | 3.43E-03 | 1.78E-03 |
| **62** | *8* | *15* | HEK293T | LMNB1 | 100 | **light fixation**, poor batch | pA | 7 | Hia5 | 200 | N | RT | B* | 500 | 52,734 | 315,838,720 | 5,989 | 9.736 | 3.58E-03 | 1.48E-03 |
| **63** | *8* | *17* | Hap1 | LMNB1 | 100 | primary at RT, poor batch | pA | 7 | Hia5 | 200 | N | RT | B* | 500 | 54,120 | 354,171,388 | 6,544 | 8.120 | 3.29E-03 | 1.72E-03 |
| **64** | *8* | *19* | HEK293T | LMNB1 | 100 | **conA beads**, poor batch | pA | 7 | Hia5 | 200 | N | RT | B* | 500 | 61,614 | 494,542,424 | 8,026 | 8.561 | 3.29E-03 | 1.41E-03 |
| **65** | *8* | *20* | GM12878 | LMNB1 | 100 | **conA beads**, poor batch | pA | 7 | Hia5 | 200 | N | RT | B* | 500 | 64,420 | 461,182,886 | 7,159 | 5.146 | 3.08E-03 | 1.52E-03 |
| **66** | *8* | *24* | Hap1 | LMNB1 | 100 | **conA beads**, poor batch | pA | 7 | Hia5 | 200 | N | RT | B* | 500 | 54,354 | 404,350,726 | 7,439 | 4.455 | 3.01E-03 | 1.65E-03 |
| **67** | *9* | *1* | HEK293T | LMNB1 | 100 | | pA | 7 | Hia5 | 200 | N | RT | B* | 500 | 45,503 | 360,723,378 | 7,927 | 10.651 | 1.84E-03 | 7.29E-04 |
| **68** | *9* | *4* | HEK293T | LMNB1 | 100 | | pA | 26 | Hia5 | 527 | N | RT | B* | 500 | 62,799 | 407,120,248 | 6,483 | 16.014 | 3.81E-03 | 1.55E-03 |
| **69** | *9* | *5* | HEK293T | LMNB1 | 100 | | pA | 7 | Hia5 | 200 | N | RT | B* | 500 | 33,562 | 297,569,878 | 8,866 | 18.086 | 3.28E-03 | 1.23E-03 |
| **70** | *9* | *6* | HEK293T | LMNB1 | 100 | | pA | 26 | Hia5 | 200 | N | RT | B* | 500 | 27,725 | 284,031,573 | 10,245 | 19.494 | 3.78E-03 | 1.42E-03 |
| **71** | *9* | *7* | HEK293T | LMNB1 | 100 | | pAG | 7 | Hia5 | 200 | N | RT | B* | 500 | 31,665 | 307,111,011 | 9,699 | 11.343 | 1.75E-03 | 6.91E-04 |
| **72** | *9* | *8* | HEK293T | LMNB1 | 50 | | pAG | 7 | Hia5 | 200 | N | RT | B* | 500 | 40,276 | 349,442,835 | 8,676 | 17.398 | 1.94E-03 | 7.89E-04 |
| **73** | *10* | *14* | HEK293T | LMNB1 | 100 | | pA | 7 | Hia5 | 200 | N | RT | B* | 500 | 202,447 | 1,902,202,299 | 9,396 | 19.705 | 2.52E-03 | 9.67E-04 |
| **74** | *10* | *15* | HEK293T | LMNB1 | 100 | RNAse | pA | 7 | Hia5 | 200 | N | RT | B* | 500 | 376,220 | 2,775,298,200 | 7,377 | 13.630 | 1.64E-03 | 6.31E-04 |
| **75** | *10* | *17* | HEK293T | LMNB1 | 100 | 300mM NaCl final wash | pA | 26 | Hia5 | 200 | N | RT | B* | 500 | 226,875 | 1,978,617,616 | 8,721 | 17.773 | 2.73E-03 | 1.13E-03 |
| **76** | *11* | *20* | HEK293T | LMNB1 | 100 | | pA | 7 | Hia5 | 200 | N | RT | B* | 500 | 129,920 | 1,273,519,587 | 9,802 | 24.331 | 2.85E-03 | 1.04E-03 |
| **77** | *11* | *21* | HEK293T | LMNB1 | 100 | | pA | 7 | Hia5 | 200 | N | RT | B* | 800 | 138,407 | 1,318,014,140 | 9,523 | 34.315 | 2.90E-03 | 1.05E-03 |
| **78** | *12* | *7* | HEK293T | LMNB1 | 50 | 2M cells | pA | 7 | Hia5 | 200 | N | RT | B* | 800 | 105,855 | 977,322,733 | 9,233 | 15.731 | 3.50E-03 | 1.41E-03 |
| **79** | *12* | *8* | HEK293T | LMNB1 | 50 | 5M cells | pA | 7 | Hia5 | 200 | N | RT | B* | 800 | 80,057 | 694,123,573 | 8,670 | 15.069 | 3.97E-03 | 1.54E-03 |
| **80** | *12* | *9* | HEK293T | LMNB1 | 50 | 10M cells | pA | 7 | Hia5 | 200 | N | RT | B* | 800 | 101,673 | 861,972,532 | 8,478 | 13.248 | 2.40E-03 | 1.03E-03 |

**Supplementary Table 1. All LMNB1-directed conditions tested.** Conditions are specified on the left side of the table, and outputs are summarized on the right hand side. For each protocol parameter (columns), the option that was selected for the final protocol is highlighted in green. ON:OFF represents the ratio of the proportion of adenines (q>=10) methylated (p>0.9) in cLADs to the proportion of adenines methylated in ciLADs, and cells are colored by the magnitude of this ratio (red=low, green=high). The neighboring columns are shaded white to blue to correspond to the proportion of adenines methylated in cLADs or across all reads. Each batch was run on a separate day. A list of abbreviations follows. Ab: Primary Antibody, Ab2: Secondary Antibody (either Goat if not specified or GP for guinea pig), Ab. Dil.: the dilution factor of the antibody (50 = 1:50), BC: barcode, RT: Room Temperature, Buffer A: same as wash buffer in final protocol, Buffer A*: same as A but with 75 mM NaCl, Buffer B: activation (methylation step) buffer in final protocol without BSA, Buffer B*: activation buffer in final protocol with BSA, conA: concanavalin A, MTase: methyltransferase, RNAse: the nuclei were treated with RNAse prior to antibody binding, SAM: S-adenosylmethionine (methyl donor), SRE XL: the Circulomics Short Read Eliminator XL kit was used to select longer fragments prior to sequencing.

| ID | Batch | BC | Cell Line | Ab | Ab dil. | Other / Notes | pA/G | Link. len. (aa) | MTase | [MTase] (nM) | Ab2 | pA/G bind temp | Act. buf. | Act. [SAM] (uM) | Read number | Total bases sequenced | Mean read len. | ON:OFF | ON-target prop. mA | All reads prop. mA |
|---|---|---|---|---|---|---|---|---|---|---|---|---|---|---|---|---|---|---|---|---|
| **81** | *4* | *22* | HEK293T | LMNB1 | 100 | **no SAM** | pAG | 29 | EcoGII | 527 | N | 4C | A | **0** | 144,251 | 1,297,558,669 | 8,995 | 1.216 | 3.83E-05 | 3.69E-05 |
| **82** | *8* | *14* | HEK293T | LMNB1 | 100 | **no SAM**, poor batch | pA | 7 | Hia5 | 200 | N | RT | B* | **0** | 37,978 | 301,486,227 | 7,938 | 2.585 | 5.13E-04 | 3.57E-04 |
| **83** | *1* | *7* | HEK293T | **IgG** | 500 | SRE XL | pAG | 29 | EcoGII | 50 | N | 4C | A | 500 | 39,167 | 872,653,096 | 22,280 | 1.113 | 2.22E-05 | 2.46E-05 |
| **84** | *2* | *13* | HEK293T | **IgG** | 500 | SRE XL | pAG | 29 | EcoGII | 150 | N | 4C | A | 500 | 66,878 | 1,500,527,829 | 22,437 | 0.982 | 4.86E-05 | 4.68E-05 |
| **85** | *4* | *2* | HEK293T | **IgG** | 100 | | pAG | 29 | EcoGII | 527 | Y | 4C | A | 500 | 165,186 | 1,608,748,957 | 9,739 | 0.818 | 6.05E-05 | 8.06E-05 |
| **86** | *4* | *3* | HEK293T | **IgG** | 100 | | pAG | 29 | EcoGII | 527 | N | 4C | A | 500 | 95,567 | 982,777,929 | 10,284 | 0.835 | 7.71E-05 | 9.15E-05 |
| **87** | *8* | *10* | GM12878 | **IgG** | 100 | | pA | 7 | Hia5 | 200 | N | RT | B* | 500 | 35,649 | 304,927,237 | 8,554 | 1.773 | 4.57E-04 | 3.72E-04 |
| **88** | *8* | *11* | HG002 | **IgG** | 100 | | pA | 7 | Hia5 | 200 | N | RT | B* | 500 | 58,023 | 418,761,436 | 7,217 | 1.448 | 4.44E-04 | 3.67E-04 |
| **89** | *8* | *12* | Hap1 | **IgG** | 100 | | pA | 7 | Hia5 | 200 | N | RT | B* | 500 | 73,437 | 453,120,139 | 6,170 | 1.865 | 4.77E-04 | 3.78E-04 |
| **90** | *8* | *13* | HEK293T | **IgG** | 100 | | pA | 7 | Hia5 | 200 | N | RT | B* | 500 | 50,184 | 395,339,186 | 7,878 | 1.563 | 4.15E-04 | 3.44E-04 |
| **91** | *12* | *16* | GM12878 | **IgG** | 50 | | pA | 7 | Hia5 | 200 | N | RT | B* | 800 | 169,811 | 1,344,179,912 | 7,916 | 1.473 | 1.50E-04 | 1.16E-04 |
| **92** | *5* | *13* | HEK293T | - | - | **free floating EcoGII** | pAG | 29 | EcoGII | 527 | - | - | A | 500 | 102,650 | 983,478,388 | 9,581 | 0.572 | 2.80E-04 | 4.49E-04 |
| **93** | *5* | *20* | HEK293T | - | - | **light fix., free floating EcoGII** | pAG | 29 | EcoGII | 527 | - | - | A | 500 | 118,802 | 260,574,348 | 2,193 | 0.573 | 8.96E-05 | 1.38E-04 |
| **94** | *8* | *16* | HEK293T | - | - | **free floating Hia5** | - | - | Hia5 | 200 | - | | B* | 500 | 49,149 | 363,924,351 | 7,405 | 1.102 | 7.55E-03 | 7.47E-03 |
| **95** | *10* | *21* | HEK293T | - | - | **free floating Hia5** | - | - | Hia5 | 200 | - | | B* | 500 | 224,739 | 1,954,375,136 | 8,696 | 1.168 | 6.34E-03 | 6.06E-03 |
| **96** | *10* | *22* | HEK293T | - | - | **free floating Hia5** w/ RNAse | - | - | Hia5 | 200 | - | | B* | 500 | 192,453 | 1,755,187,759 | 9,120 | 1.228 | 6.96E-03 | 6.45E-03 |
| **97** | *12* | *17* | GM12878 | - | - | **free floating Hia5** | - | - | Hia5 | 200 | - | | B* | 800 | 94,126 | 831,589,618 | 8,835 | 1.123 | 7.98E-03 | 7.66E-03 |

**Supplementary Table 2. Control conditions tested.** Same as Supplementary Table 1 but for 3 different negative control conditions (not expected to show methylation, or not expected to show enrichment in cLADs): no SAM added at activation/methylation step, nonspecific IgG isotype control antibody used, or free-floating enzyme was added at the activation/methylation step to methylate all accessible DNA.

| Sample number | Target / description | Cell line | Reads | Bases | Mean read length |
|---|---|---|---|---|---|
| 1 | CTCF | GM12878 | 729097 | 8824849107 | 12104 |
| 2 | CTCF | GM12878 | 209699 | 2864997263 | 13662 |
| 3 | CTCF | GM12878 | 215664 | 7052382103 | 32701 |
| 4 | free pA-Hia5 | GM12878 | 970282 | 10371517658 | 10689 |
| 5 | IgG | GM12878 | 1421526 | 11531597797 | 8112 |
| 6 | H3K9me3 | HG002 | 896511 | 8057506035 | 8988 |
| 7 | H3K9me3 | HG002 | 155656 | 1847831908 | 11871 |
| 8 | H3K9me3 | HG002 | 233920 | 5798858000 | 24790 |
| 9 | free pA-Hia5 | HG002 | 71713 | 1433794520 | 19994 |
| 10 | free pA-Hia5 | HG002 | 204504 | 3771902818 | 18444 |
| 11 | free pA-Hia5 | HG002 | 64758 | 1219166120 | 18826 |
| 12 | free pA-Hia5 | HG002 | 145380 | 2878390589 | 19799 |
| 13 | IgG | HG002 | 306270 | 7857066233 | 25654 |
| 14 | IgG | HG002 | 132045 | 2685488328 | 20338 |
| 15 | in vitro methylated genomic DNA | GM12878 | 330573 | 2785142705 | 8425 |
| 16 | unmethylated genomic DNA | GM12878 | 437135 | 4040533340 | 9243 |
| 17 | CENP-A | HG002 | 258948 | 2278497590 | 8799 |
| 18 | CENP-A | HG002 | 325860 | 2490299385 | 7642 |
| 19 | CENP-A | HG002 | 65205 | 454714846 | 6974 |
| 20 | free pA-Hia5 | HG002 | 230216 | 2018755168 | 8769 |
| 21 | free pA-Hia5 | HG002 | 114364 | 848652995 | 7421 |
| 22 | free pA-Hia5 | HG002 | 257095 | 1740051056 | 6768 |
| 23 | IgG | HG002 | 252694 | 2349991082 | 9300 |
| 24 | IgG | HG002 | 235264 | 2068374069 | 8792 |
| 25 | IgG | HG002 | 84856 | 624068929 | 7354 |
| 26 | untreated | HG002 | 260016 | 2167665966 | 8337 |
| 27 | untreated | HG002 | 436472 | 3262286345 | 7474 |
| 28 | untreated | HG002 | 86613 | 573212999 | 6618 |

**Supplementary Table 3. Sequencing summary metrics.** The number of reads, bases, and mean read length are indicated for CTCF-, H3K9me3-, and CENP-A-directed DiMeLo-seq, along with accompanying controls.
